# Molecular mechanism of light-driven sodium pumping

**DOI:** 10.1101/2020.01.29.925347

**Authors:** K. Kovalev, R. Astashkin, I. Gushchin, P. Orekhov, D. Volkov, E. Zinovev, E. Marin, M. Rulev, A. Alekseev, A. Royant, P. Carpentier, S. Vaganova, C. Baeken, I. Sergeev, D. Zabelskii, T. Balandin, G. Bourenkov, V. Borshchevskiy, G. Büldt, E. Bamberg, V. Gordeliy

## Abstract

Microbial rhodopsins appeared to be the most abundant light-harvesting proteins on the Earth and are the major contributes to the solar energy captured in the sea. They possess highly diverse biological functions. Explosion of research on microbial rhodopsins led to breakthroughs in their applications, in particular, in neuroscience.

An unexpected new discovery was a Na^+^-pumping KR2 rhodopsin from *Krokinobacter eikastus*, the first light-driven non-proton cation pump. A fundamental difference between proton and other cation pumps is that non-proton pumps cannot use tunneling or Grotthuss mechanism for the ion translocation and, therefore, Na^+^ pumping cannot be understood in the framework of classical proton pump, like bacteriorhodopsin. Extensive studies on the molecular mechanism of KR2 failed to reveal mechanism of pumping. The existing high-resolution structures relate only to the ground state of the protein and revealed no Na^+^ inside the protein, which is unusual for active ion transporters.

KR2 is only known non proton cation transporter with demonstrated remarkable potential for optogenetic applications and, therefore, elucidation of the mechanism of cation transport is important. To understand conception of cation pumping we solved crystal structures of the functionally key O-intermediate state of physiologically relevant pentameric form of KR2 and its D116N and H30A key mutants at high resolution and performed additional functional studies.

The structure of the O-state reveals a sodium ion near the retinal Schiff base coordinated by N112 and D116 residues of the characteristic (for the whole family) NDQ triad. The structural and functional data show that cation uptake and release are driven by a switching mechanism. Surprisely, Na^+^ pathway in KR2 is lined with the chain of polar pores/cavities, similarly to the channelrhodopsin-2. Using Parinello fast molecular dynamics approach we obtained a molecular movie of a probable ion release.

Our data provides insight into the mechanism of a non-proton cation light-driven pumping, strongly suggest close relation of sodium pumps to channel rhodopsins and, we believe, expand the present knowledge of rhodopsin world. Certainly they might be used for engineering of cation pumps and ion channels for optogenetics.

---

Microbial rhodopsins (MRs) are transmembrane light-sensitive proteins, found in archaea, bacteria, eukaryotes and also viruses^1^. They possess diverse biological functions and are the core of breakthrough biotechnological applications, such as optogenetics. MRs are composed of seven transmembrane α helices (A-G) with the cofactor retinal covalently bound to the lysine residue of helix G via the Schiff base (RSB). Due to a conflict of a presence of a cation close to the RSB proton, it was believed that Na^+^-pumping rhodopsins could not exist in nature. Despite this paradigm, the first light-driven Na^+^ pump KR2 was identified in *Krokinobacter eikastus* in 2013^2^. Its functional and structural properties were extensively studied^3–5^. KR2 contains a characteristic for all known Na^+^-pumping rhodopsins (NaRs) set of N112, D116, Q123 residues in the helix C (NDQ motif). It was shown that the protein pumps Na^+^ when its concentration is much higher than that of H^+^, which is characteristic for physiological conditions, otherwise it acts as a H^+^ pump^6^. An extensive mutational analysis of KR2 indicated key functional residues, such as N112, D116, Q123, but also H30, S70, R109, R243, D251, S254, G263^2, 3, 7, 8^. Moreover, potassium-pumping and potassium-channeling variants of KR2 were designed, making the protein a potential tool for optogenetics^3–5, 9^.

A key question remains to be answered: what is the mechanism of pumping. Indeed, principles of Na^+^ transport by KR2 and other NaRs remains unclear. After light excitation the photocycle starts with the retinal isomerization from all-*trans* to 13-*cis* configuration^13^. In the Na^+^-pumping mode the protein is characterized by the K, L/M and O intermediates^13^ (Fig. 1A). It is known that upon retinal isomerization the proton is translocated from the Schiff base (RSB) to the D116 during M-state formation^2, 14^. Uptake of sodium occurs in the M-to-O transition. It is hypothesized that Na^+^ may pass the cytoplasmic gate comprised by Q123 and the neutralized RSB, and binds in the central region near D116, N112 and presumably D251 transiently in the O-state^4, 11, 13^. It was also suggested that with the decay of the O-state, Na^+^ is released via R109 and the cluster of E11, E160 and R243 to the extracellular space^3, 5^. Therefore, the O-state is considered to be the key for elucidation of the Na^+^-pumping mechanism^13^.

**Figure 1.**
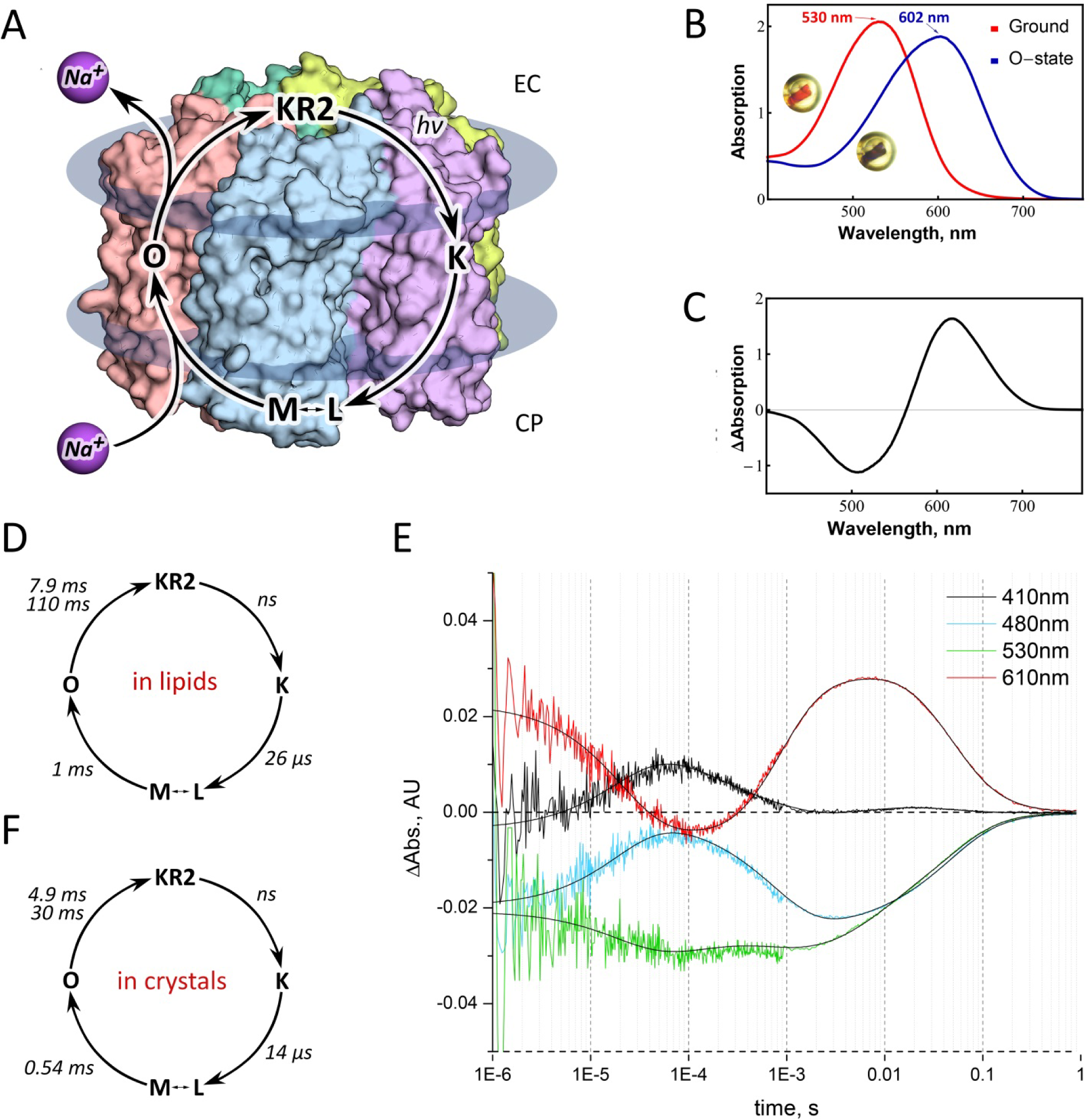
Spectroscopy of KR2 in crystals. **A.** Scheme of the KR2 photocycle indicates that Na^+^ binding occurs transiently in the red-shifted O-state. **B.** UV-visible absorption spectra measured *in crystallo* at 100 K of the Ground state (red) and the O-state of KR2 (insets: photos of the same frozen KR2 crystal in the cryoloop before and after laser illumination, near the corresponding spectra). **C.** Difference spectrum calculated between the blue and red spectra shown in Fig. 1B. **D.** Photocycle of KR2 reconstituted in DOPC^2^. **E.** Time traces of absorption changes of KR2 microcrystals at 410 (black), 480 (light blue), 530 (green) and 610u (red) probe wavelengths. Black lines indicate fitting lines based on the sequential kinetic model shown in **F.** Photocycle of KR2 in microcrystals, determined in the present work.

It is important that the protein always forms pentamers being reconstituted into lipid membrane^10^. KR2 pentamers appear also under physiological conditions in crystals^3, 4, 11^ and detergent micelles^4, 12^. Hence, KR2 is considered to be a pentamer in the native membrane (Fig. 1A). Pentamerization is important for Na^+^ pumping by KR2^4^. Particularly, all previous functional investigations of KR2 were performed on the pentameric form of the protein. Moreover, it was recently reported that in the ground state under physiological conditions (PDB ID: 6REW^4^) KR2 has a large water-filled cavity near the RSB (Schiff base cavity). This conformation of the protein was called ‘expanded’^4, 11^, and only occurs in the pentameric form of KR2. On the contrary, in the monomeric form the large cavity is absent, and the protein is in another conformation, called ‘compact’^4, 11^. Thus, to elucidate the mechanism of light-driven sodium pumping one needs to study the biologically relevant pentameric form of KR2, where it forms the ‘expanded’ conformation in the ground state.

We should note that uncovering of mechanism of Na^+^ pumping is of great importance and simultaneously is a challenge. First of all, it should be remarkably different from that of H^+^ pumping^13^ and, therefore, huge amount of our knowledge on the H^+^ pumping mechanism obtained with a classic proton pump bacteriorhodopsin (BR)^15, 16^ could not be applied straightforward to NaRs. Not only there is a conflict of simultaneous presence of two positive charges in close proximity, the protonated RSB (RSBH^+^) and Na^+^ in the case of KR2, but also there are fundamental differences in the translocation of the proton and other cations. Namely, proton transport in biological objects implies ion tunneling, which is dramatically hampered in case of larger cations. Furthermore, non-proton cation pumps cannot utilize the Grotthuss mechanism for ion translocation^15^. Hence, pathways of proton in light-driven pumps cannot be the same as in the cation pumps like NaRs. All history of studies of BR, halorhodopsin (HR) and also sensory rhodopsin II (SRII) shows that high-resolution structures of the intermediate states of a rhodopsin are key for the understanding of the mechanisms. It is even more valid in the case of NaRs, as Na^+^ inside the protein is absent in the ground state of KR2, which provides a wide room for speculations on the mechanism of Na^+^ transport.

Here we present the structures of the O-state of physiologically relevant pentameric form of KR2 – the key intermediate of Na^+^ pumping – and functionally important D116N and H30A mutants of the protein. The structure of the O-state reveals Na^+^ binding site inside the rhodopsin and together with the structures of the mutants allows us to elucidate key determinants of cation pumping by KR2.

## RESULTS

We crystallized KR2 in the functional state (at pH 8.0) using *in meso* approach similarly to our previous works^4, 17, 18^. To verify that the protein in crystals undergoes the same photocycle as in lipids, we performed time-resolved visible absorption spectroscopy on KR2 microcrystal slurries. The experiments showed that similar to the protein in detergent micelles and lipids, crystallized KR2 also forms characteristic K-, L/M- and O-states (Fig. 1). Then the O-state was trapped using an approach described in ref ^18^. In brief, we illuminated KR2 crystals with 532-nm laser to freeze-trap the O-state. The cryostream was blocked for 1 second during laser exposure and released back before turning off the laser. This procedure allows accumulation and trapping in crystals of the dominant intermediate of the protein photocycle^18, 19^. Single-crystal spectrophotometry showed that after the procedure nearly all proteins in the crystal were trapped in the red-shifted intermediate state (Fig. 1B, C). KR2 undergoes only two red-shifted states during photocycle: the K- and the O-states (Fig. 1F). As the difference electron density maps (described in details below) do not indicate even a low fraction of the ground state in the structure, show the Na^+^ bound near the RSB region, which is characteristic for the O-state of KR2, and the O-state is a dominant intermediate of the KR2 photocycle, we consider the trapped intermediate as solely the O-state.

To verify that flash-cooling does not affect the conformation of the O-state, we collected X-ray diffraction data at room temperature (RT) during continuous 532-nm laser illumination with single crystals of KR2. This procedure also allows to detect the dominant conformational changes of the photocycle^19^. We collected RT crystallographic data at 2.6 Å resolution by merging of 3 complete datasets obtained from 3 single crystals (see Supplementary Text) and structure refinement identified nearly 1/1 ratio of the ground/O-states populations in the crystals. The structure of the O-state at RT is identical to that at 100K, which indicates that cryo-cooling does not affect the O-state of KR2. Hence, we describe further only the structure of cryo-trapped intermediate, since it has higher resolution (2.1 Å) and occupancy of the O-state (100 %).

Using the crystals with the trapped intermediate, we solved the structure of the O-state at 2.1 Å (Supplementary Table 1). The crystal symmetry and lattice parameters are the same as described previously for the ground state of the protein, with one KR2 pentamer in the asymmetric unit^3, 4^.

The structure demonstrates notable rearrangements compared to the ground state of KR2 (Fig. 2 and Supplementary Fig 1). The root mean square deviation (RMSD) between the backbone atoms of the pentamers and protomers of the ground (PDB ID: 6REW^4^) and the O-states (present work) are 0.55 and 0.53 Å, respectively. The main changes occur in the extracellular parts of the helices B and C, which are shifted by 1.0 and 1.8 Å, respectively (Supplementary Fig 1). Helices A, D and G are also displaced by 0.7 Å in the extracellular regions (Supplementary Fig 1).

**Figure 2.**
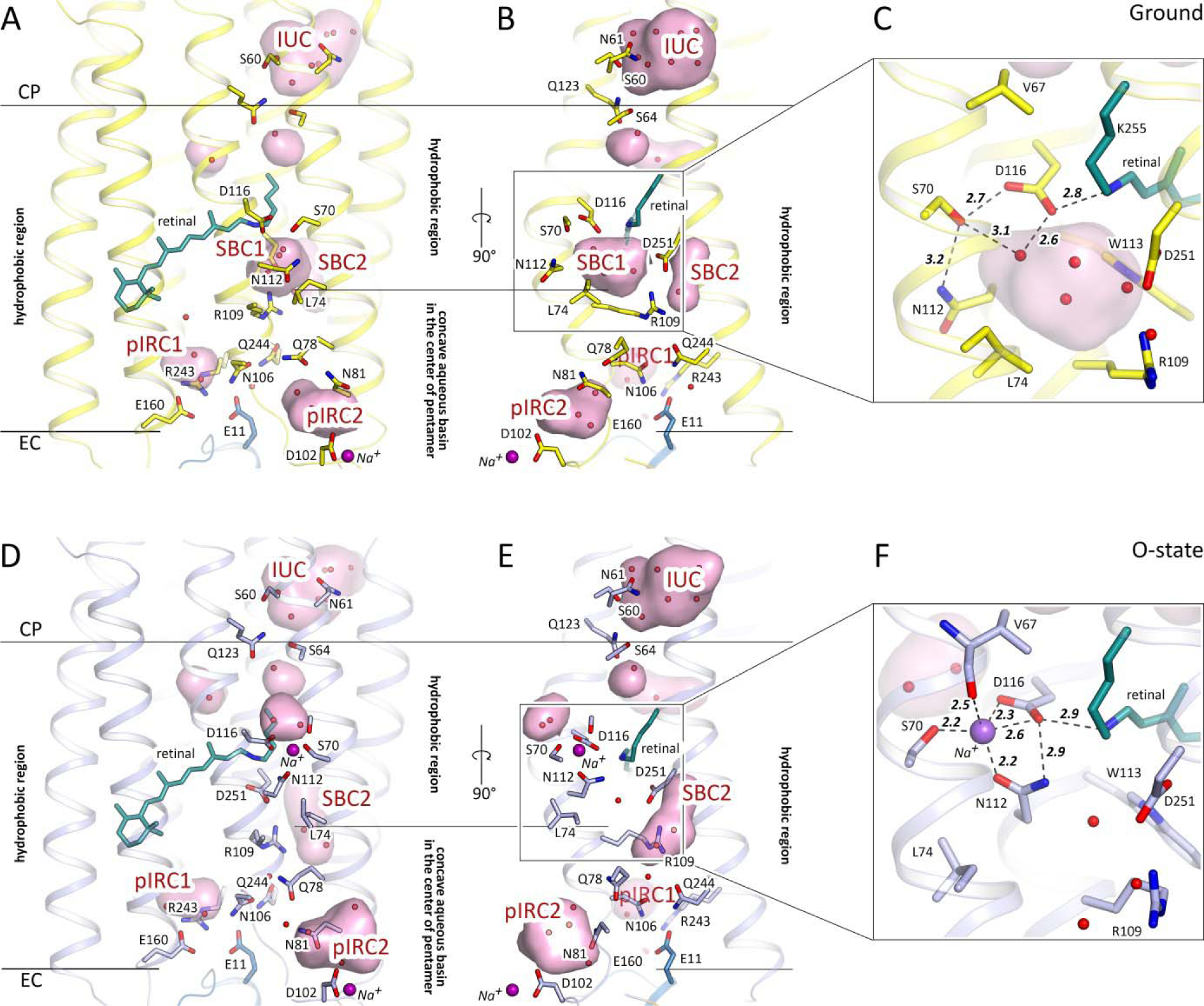
Overall comparison of the ground and O-states of KR2. **A; D.** Side view of the KR2 protomer in the ground (yellow, PDB ID: 6REW) and O- (blue, present work) states. **B, E.** View from the side of the helices A and B. Membrane hydrophobic/hydrophilic boundaries are shown with the black lines. The membrane boundary at the extracellular side is located at two levels for the inner and outer part of the KR2 pentamer respectively. Helices A and B face the concave aqueous basin, formed in the central pore of the pentamer and helices C-G face the lipid bilayer, surrounding the pentamer. Water molecules are shown as yellow and blue spheres for ground and O-state respectively. Helices A and B are hidden for clarity. **C, F.** Detailed view of the RSB region of the ground and the O-state of KR2. Cavities (ion-uptake cavity – IUC; the Schiff base cavities 1 and 2 – SBC1 and SBC2, respectively; putative ion-release cavities 1 and 2 – pIRC1 and pIRC2, respectively) inside the protein were calculated using HOLLOW^49^ shown in pink and marked with red labels. Retinal cofactor is colored teal. Water molecules are shown with red spheres. Sodium ion is shown with a purple sphere. Hydrogen bonds involving S70, N112, D116, D251 and RSB are shown with black dashed lines. The lengths of the shown hydrogen bonds are shown with bold italic numbers and are in Å. Helix A and SBC2 are hidden for clarity.

### Retinal binding pocket of KR2 in the O-state

The polder^20^ electron density maps built around the retinal cofactor strongly suggest a distortion around C14. Nevertheless, the retinal is in all-*trans* conformation in the O-state (Supplementary Fig 2). Surprisingly, this is in contrast to the recently published time-resolved Fourier-transform infrared spectroscopy (FTIR) data, where authors suggested 13-*cis* configuration in the O-state of NaRs^21^. However, it is in line with the data on another light-driven Na^+^ pump from *Gillisia limnaea* (GLR), published in 2014, where the authors report a distorted all-*trans* configuration of retinal in the O-state^22^. Such retinal configuration means that relative locations of the RSBH^+^ and D116 side chain are similar to those in the ground state (PDB ID: 6REW^4^). RSBH^+^ is hydrogen bonded to D116 and the distance between them is 2.9 Å in the O-state (Fig. 2F). The existence of this hydrogen bond is supported by time-resolved resonance Raman spectroscopy^14^. The positions of the residues comprising the retinal pocket, particularly W113, D251, D116, I150, Y218 and W215 are also shifted relative to that in the ground state (Supplementary Fig 3).

### Sodium binding site inside the protein

The crystal structure of the O-state of KR2 clearly reveals the Na^+^ binding site near the RSB, comprised of S70, N112 and D116 side chains and main chain oxygen of V67 (Fig. 2F and Supplementary Fig 2, 4). Previous mutational analysis confirms the importance of these residues for KR2 pumping activity. Indeed, D116 is crucial for KR2 functioning^2^, and N112 determines ion selectivity^7^. Substitution of S70 with threonine or alanine dramatically decreases Na^+^-pumping activity of KR2^5, 9^. The mean distance between Na^+^ and the coordinating oxygen atoms is 2.3 Å (Fig. 2F and Supplementary Fig 2).

While in the ground state KR2 is in the ‘expanded’ conformation, in the O-state N112 is flipped towards S70 and D116, therefore the overall configuration is similar to that of the ‘compact’ conformation of KR2^4, 11^ (Fig. 2, Supplementary Fig 5, 6). This is also evidenced by disappearance of the big polar cavity near the RSB (SBC1) and enlargement and elongation of the cavity near R109-D251 pair (SBC2) in the O-state (Fig. 2 and Supplementary Fig 6). Four water molecules, filling the SBC1 in the ‘expanded’ ground state (Fig. 2C), are displaced as follows: two of them are found in the small cavity formed in the intermediate near S70 at the pentamerization interface, one remains at the same place and is coordinated by N112 and D251, and the last one is moved to the SBC2 near L75 and R109 at the inner extracellular part of the protein (Fig. 2F). Upon sodium binding and formation of the ‘compact’ state L74 side chain also flips simultaneously with the N112 in order to avoid the steric conflict of these two residues. Our mutational analysis indicated that L74A substitution dramatically decreases pumping activity of the protein (Supplementary Fig 7). Hence, this additionally supports the importance of the ‘compact’ conformation for Na^+^ pumping by KR2.

Interestingly, the location of the sodium binding site in the O-state of KR2 is similar to that of the chloride ion binding sites in the ground state of halorhodopsins (Supplementary Fig 8). Namely, in a chloride-pumping rhodopsin from *Nonlabens marina* S1-08 (ClR)^23, 24^ the anion is coordinated by the N98 and T102 of the NTQ motif, which are analogous to the N112 and D116 of the NDQ motif of light-driven sodium pumps (Supplementary Fig 8).

### ‘Expanded’-to-‘compact’ and ‘compact’-to-‘expanded’ conformational switches are key determinants of Na^+^ uptake and release

The similarity of protein conformation in the O-state to the ‘compact’ is intriguing, however, could easily be explained. Indeed, relative location of the RSBH^+^ and Na^+^-D116^-^ pair makes the distribution of the charges in the central part of the protein nearly identical to that of the KR2 with protonated D116 at acidic pH^2–4^. The structures of KR2 in the ‘compact’ conformation are also observed only at low pH^3–5^. It was thus suggested that the ‘compact’ conformation may appear in response to the D116 neutralization^4^. To understand better the nature of conformational switches in KR2 and the influence of D116 protonation on the protein conformation, we produced and crystallized KR2-D116N at pH 8.0, which mimics the WT protein with fully protonated D116 and solved its structure in the pentameric form at 2.35 Å.

Confirming our hypothesis, the structure shows that introduction of asparagine at the position of D116 led to the flip of the side chains of N112 and L74 in comparison to the ground state of the wild type (WT) protein and disappearance of the SBC1, characteristic for the ‘expanded’ conformation (Supplementary Fig 6). Overall, the structure of D116N is very similar to that of the O-state (RMSD 0.2 Å) and also to the ‘compact’ conformation (RMSD 0.3 Å) of the KR2-WT (Supplementary Fig 6). However, Na^+^ is absent inside the protomers and the relative orientation of the RSBH^+^ and N116 is altered (Supplementary Fig 6). Particularly, the RSBH^+^ forms two alternative conformations and hydrogen bond between the RSBH^+^ and N116 is absent (see Supplementary Text).We also observed that D116 protonation destabilizes the pentameric assembly of KR2 (see Supplementary Text). Thus, we suggest that neutralization of D116 is the key determinant of the formation of the ‘compact’ conformation, which explains their structural similarity.

The structures of KR2 O-state and D116N mutant, together with previously described pH dependence of the KR2 organization^4^, allow us to conclude that the ‘compact’ conformation and, particularly, N112 flip towards D116, stabilizes neutralized RSB counterion and correspondingly neutral transiently formed Na^+^-D116^-^ pair during the photocycle.

It was suggested previously that the SBC1 in the ‘expanded’ conformation surrounded by R109, N112, W113, D116 and D251 might be a transient Na^+^ binding site in an intermediate state of the protein photocycle^4, 11^. However, the present work shows that Na^+^ binds far from R109 and D251 (Fig. 2F). Since the release of Na^+^ occurs upon the O-to-ground state transition, which structurally corresponds to ‘compact’-to-‘expanded’ switch, we suggest that the ‘expanded’ conformation is also important for the ion release to the extracellular space. Importantly, Na^+^ uptake and release are guided by the switch from the ‘expanded’ to the ‘compact’ and then again back to the ‘expanded’ conformations, respectively.

### Sodium translocation pathway

Although the structure of KR2 protomer near the RSB is altered in the O-state, the organization of both putative ion uptake and ion release regions remains the same to those in the ground state (Fig. 3). It is not surprising when we consider the cytoplasmic part of the protein. In the ground state Na^+^ does not penetrate to the inside of KR2. At the same time, in the O-state Na^+^ should not have a way to return back to the cytoplasm. Therefore, the pathway, formed upon transition from the ground to the O-state, connecting the ion uptake cavity (IUC) with the RSB region should be blocked in both ground and the O-states. Consequently, the restoration in the O-state of the initial conformation of S64-Q123 pair, separating the IUC from the RSB environment, is expected. On the other hand, the same organization of the E11-E160-R243 cluster and putative ion release cavity (pIRC1) in the ground and O-states may seem quite surprising. Previously, these residues were suggested to line the pathway of Na^+^ release to the extracellular bulk during the direct O-to-ground transition^3–5^. The absence of the disturbance in this region means either that the energy stored in the distorted all-*trans* retinal in the O-state is enough to relocate Na^+^ directly from the core of the protein to the bulk without any transient binding sites or/and that there might be another ion release pathway in KR2. Indeed, the second hypothesis is supported by the mutational analysis, which showed that substitution of E11, E160 or R243 to alanines or polar non-charged residues does not abolish Na^+^-pumping activity, however, affects the stability of the proteins^2, 3, 5^. The other (alternative) putative way for Na^+^ release goes from the inner extracellular part of the protein through the elongated in the O-state SBC2 to the bulk near the Na^+^ bound at the surface of KR2 in both the ground and O-states. These channel-like pathway is constricted with the only side chain of Q78 residue (Fig. 2, 3D). The pathway propagates from the inner region between Q78, N106 and R109 to the relatively large cavity (pIRC2) between helices B and C, BC loop and helix A’ of adjacent protomer at the extracellular side (Fig. 2, 3 and Supplementary Fig 9). The cavity proceeds further to a concave aqueous basin facing the extracellular solution, formed in the central pore of the KR2 pentamer at the extracellular side and is surrounded by Q78, N81, S85, D102, Y108 and Q26’ residues and filled with water molecules in both the ground and O-states (Fig. 3). Notable displacements of these residues and waters occur in the O-state, such as the flip of N81 towards H30’ of the adjacent protomer (Supplementary Fig 10). Consequently, in the O-state additional water molecule appears in the pIRC2, which is coordinated by hydrogen bonds with E26’, H30’ and N81 (Supplementary Fig 10). The positions of Q78 and Y108 are also altered in the O-state (Supplementary Fig 10).

**Figure 3.**
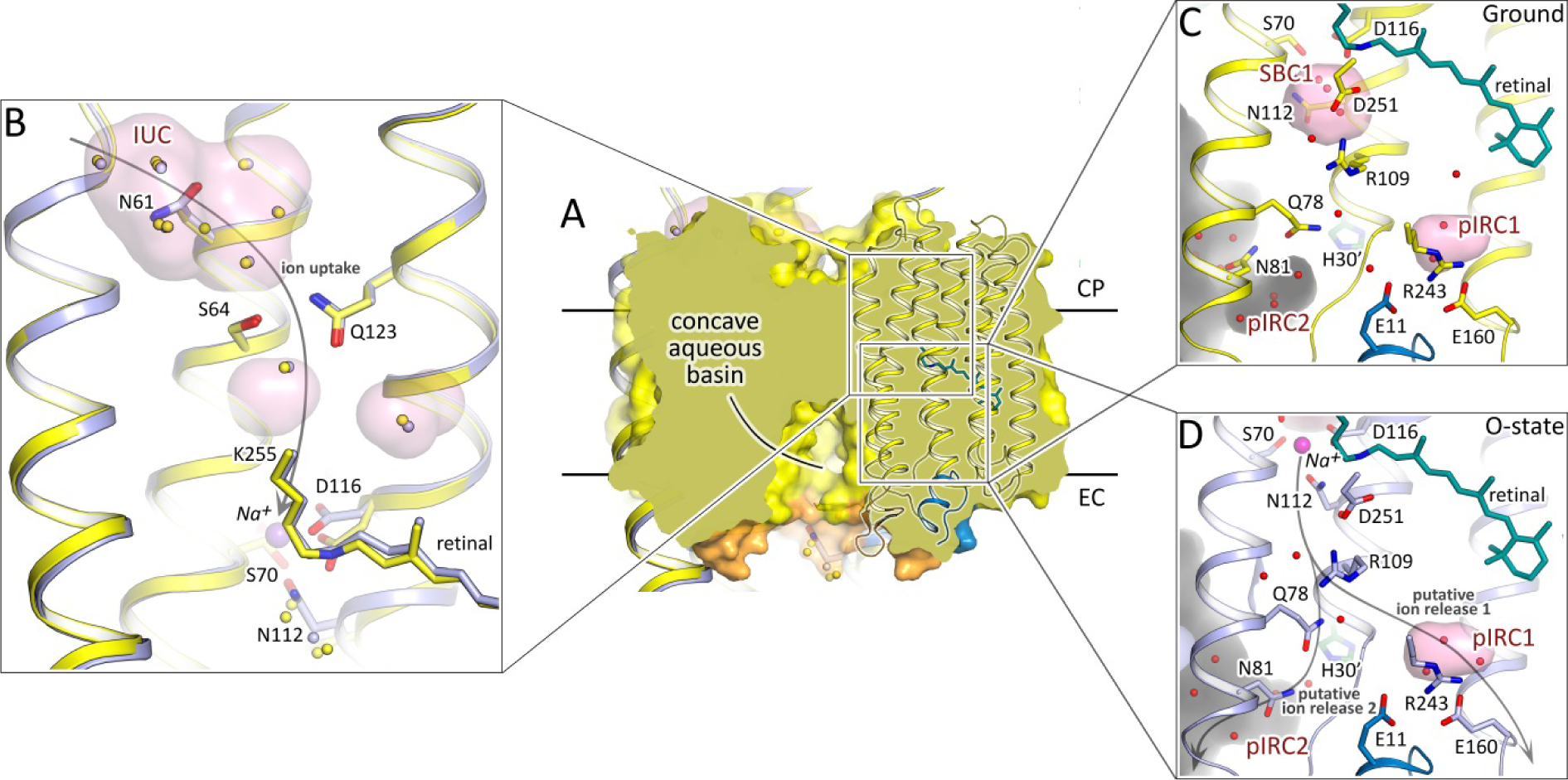
Ion uptake and release pathways of KR2. **A.** Section view of KR2 pentamer in the membrane. Concave aqueous basin facing the extracellular space is indicated by the black line. Only one protomer is shown in cartoon representation. Membrane core boundaries are shown with black lines. **B.** Structural alignment of the cytoplasmic parts of the ground (yellow) and O-(blue) states of KR2. Water molecules are shown with yellow and blue spheres for the ground and O-state, respectively. **C.** Detailed view of the extracellular side of KR2 in the ground state. **D.** Detailed view of the extracellular side of KR2 in the O-state. Cavities inside the protein are calculated using HOLLOW^49^ shown in pink and marked with red labels. Protein surface concavity from the aqueous basin at the extracellular side is colored gray. Retinal cofactor is colored teal. Water molecules are shown with red spheres. Sodium ion is shown with purple sphere. N-terminal α elix is colored blue. BC loop is colored orange. H30’ of adjacent protomer lored with dark-green. Helices A, F and G are hidden for clarity. Gray arrows identify putative ion uptake and two ion release pathways.

To probe the possible ion release pathways, we conducted 10 short molecular dynamics simulations starting from the sodium-bound conformation (Supplementary Fig 11 and Supplementary Vid. 1). The simulations revealed that the R109 sidechain forms a barrier for sodium exiting SCB1, and changes its position to allow sodium passage. Upon passing R109, Na^+^ either exits the protein via pIRC2 (8 simulations out of 10), or proceeds towards pIRC1 (2 simulations). In the first scenario, the ion is quickly released towards the aqueous basin in the middle of KR2 pentamer in the vicinity of another ion found at the interface between the protomers, and is sometimes observed to replace it (Supplementary Vid. 1). In the second scenario, the ion samples different locations around the E11-E160-R243 triad and is later released via Asn106 and Gln157 on the outer side of the pentamer.

Importantly, the organization of the pIRC2 region is identical in the O-state and KR2-D116N (Supplementary Fig 12A). It means that the rearrangements on the surface of the KR2 occur not directly in response to the retinal isomerization upon photon absorption, but rather due to the redistribution of charges in the central inner part of the protein protomer. Such long-distance interactions between the RSB-counterion and pentamer surface were already studied for the WT and H30A variant of KR2^12, 25^. To gather more details about these interactions, we solved the structure of pentameric form of KR2-H30A at pH 8.0 at 2.2 Å.

Overall, the structure of this mutant is nearly the same as that of the ground state of WT protein (RMSD 0.15 Å) (Supplementary Fig 5, 13). The organization of their inner cytoplasmic, central and extracellular parts is identical. Surprisingly, in contradiction to the earlier FTIR experiments, Na^+^ bound at the oligomerization interface of the WT protein is also present in H30A^2^. However, the region of H30 is altered (Supplementary Fig 12). Particularly, H30 side chain is replaced by two additional water molecules (w_B_’ and w_B_’’) (Supplementary Fig 12). Moreover, the H30A mutation leads to appearance of second alternative conformation of Y108, not identified in other structures of KR2 or its variants (Supplementary Fig 12). It was shown that H30A is more selective to sodium and almost does not pump protons^2^. As the only existing differences in the structures of the WT and the mutant occur near Q78, N81, Y108, H30’ and pIRC2, we suggest that this region is important for cation selectivity, which additionally support the hypothesis that this region is a part of ion translocation pathway.

To verify the suggested ion-release pathway we performed functional studies of KR2 mutants in *E.coli* cells suspension, similar to previous works^3, 26^ (Supplementary Fig 7). The results showed that Q78 is the key residue, which is likely to act as a gate for sodium, flipping upon sodium passage (Q78L mutant remains almost fully functional). The blocking of Q78 motion (Q78Y,W mutations) resulted in dramatic decrease of the pumping activity. In Q78A mutant, similar to Q123A, the sodium pumping activity is retained, however decreased notably in comparison to the WT protein. Another interesting finding was that Y108A mutant almost fully lost its pumping ability.

Hence, we suggest that Na^+^ translocation pathway propagates from IUC to pIRC2 via a chain of polar inner cavities, which are modified during photocycle. IUC and pIRC2 are separated from the inside of KR2 by two weak gates near Q123 and Q78, respectively. This makes the KR2 ion pathway similar to that of the channelrhodopsin 2^27^ (*Cr*ChR2) (Supplementary Fig 14). However, unlike in CrChR2, KR2 has the Na^+^ binding site in the central region near the RSB.

**Figure 4.**
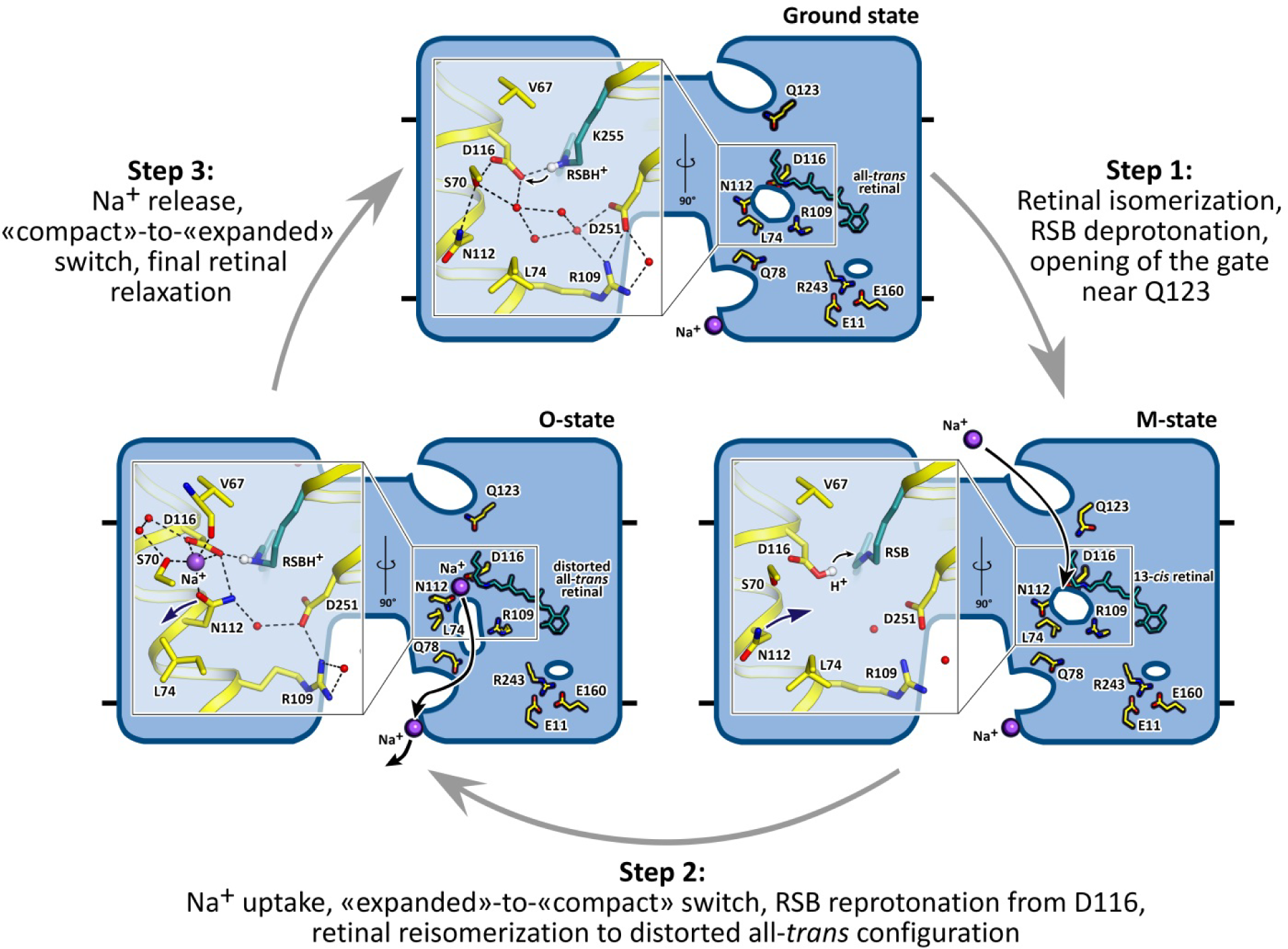
Proposed Na^+^ pumping mechanism. Schematic side section view of the KR2 pentamer is shown. Membrane core boundaries are shown with black lines. Cavities are demonstrated as white ellipses. Enlarged view of the RSB region is shown at the left part of the pentamer. The 13-*cis* configuration of the retinal cofactor is modelled manually for schematic representation. Na^+^ is shown with violet spheres. Black arrows indicate proposed Na^+^ uptake and release pathways. Violet arrows indicate rearrangement of N112 side chain during “expanded”-to-“compact” and back “compact”-to-“expanded” switches. Small gray arrows indicate the translocation of the hydrogen from the Schiff base to D116 during the formation of the M-state and following reprotonation of the Schiff base from the D116 in the M-to-O transition. Retinal cofactor is colored teal. Waters are shown with red spheres. Hydrogen bonds in the RSB region are shown with black dashed lines.

### Mechanism of sodium pumping

Crystal structures of the O-state and D116N and H30A mutants of KR2 together with available literature data allow us to suggest a mechanism of protein functioning (Fig. 4).

*Step 1.* In the ground state KR2 is in the ‘expanded’ conformation with the SBC1 filled with four water molecules. The RSBH^+^ is hydrogen bonded to its counterion D116^-^. The Q123-S64 gate separates the IUC and the RSB. With the absorption of the light photon the retinal isomerizes from all-*trans* to 13-*cis* configuration and the red-shifted K-state appears in nanoseconds, followed by the formation of the L/M intermediate in about 30 μ. The proton is translocated from the RSBH^+^ to the D116^-^ with the formation of the M-state and the hydrogen bond between them is absent in this intermediate. The Q123-S64 gate also opens in this step.

*Step 2.* With the rise of the O-state Na^+^ passes the gate and deprotonated RSB and binds between S70, N112 and D116 belonging to helices B and C. Na^+^ uptake results in the proton translocation from D116 back to the RSB with the restoration of the RSB-counterion hydrogen bond, thus preventing the Na^+^ backflow to the cytoplasmic side. It also causes the flip of N112 side chain for the stabilization of Na^+^-D116^-^ pair, therefore, the ‘compact’ state appear at this step. Retinal is still in the distorted all-*trans* conformation in the O-state.

*Step 3.* With the decay of the O-state retinal returns completely to its ground configuration and the ‘expanded’ conformation of KR2 occurs with the N112 flip back to the pentamerization interface, thus opening the way for Na^+^ release. In the O-state the release pathway is prepared as it is evidenced from the O-state structure. We suggest that the release preferentially involves Q78 and proceeds towards the cavity, formed by N81, Y108 and H30’ of adjacent protomer (pIRC2). We also suggest that four water molecules, filling the SBC1 in the dark state are involved in the Na^+^ hydration during its transitions inside the protein. The Na^+^ release might proceed using the relay mechanism, when the ion, released from the KR2 protomer, replaces the ion, bound at the protein oligomerization interface. The proposed relay mechanism allows lowering the energy barriers for facilitation of the Na^+^ release. The E11-E160-R243 cluster and the cavity near it (pIRC1), suggested earlier to be involved in Na^+^ release, thus may be involved mainly in protein stabilization, rather than in ion translocation pathway.

Last but not least, in the absence of Na^+^ KR2 acts as a proton pump with the significantly altered photocycle and the absence of the pronounced O-state. The long living L/M/O-like state decays slowly in the proton-pumping mode^2^. We note that this is in agreement with the suggested mechanism of Na^+^ pumping. We suggest that the formation of the K-, L- and M-states is the same for the Na^+^- and H^+^-pumping modes, however, in the absence of Na^+^ the ion does not flow diffusely to the central region when the RSB is neutral, and RSB is presumably reprotonated from the cytoplasmic side through the IUC and no rearrangements occur in the region of D116. Then the slow relaxation of the retinal to all-*trans* configuration triggers the proton release from the D116 to the extracellular side and the protein returns to the ground state.

## SUMMARY

The presented here structure of the key functional O-intermediate state of the KR2 rhodopsin allowed us to propose the pathway and molecular mechanism of the active light-driven Na^+^ transport. We suggest that the pathway of Na^+^ connects two aqueous concavities in the cytoplasmic and extracellular parts of the KR2, respectively, via several polar cavities inside the protein, which are separated from each other by weak gates. From that side, the organization is very similar to that of *Cr*ChR2^27^. The principal difference is that in opposite to *Cr*ChR2, KR2 has the tight transient Na^+^ binding site near the Schiff base, separating the cytoplasmic and + extracellular parts of the protein. Judging from the arrangement of the cavities in the O-state, Na transport in KR2 appears rather like in a perfectly outwardly directed Na^+^ channel than in a pump, which we interpret as a chimera between the two fundamental mechanisms. Similar concept was suggested earlier^13^, however, it mostly concerned the diffusion Na^+^ uptake from the cytoplasmic part. The presence of the positively charged R109 side chain at the extracellular part of the protein presumably prevents Na^+^ uptake from that side in the M-state, as also evidenced by a recent finding that R109Q mutation converts KR2 into a potassium channel^9^. Presented here model of the O-state together with molecular dynamics simulations and mutational analysis, as well as the absence of any intermediate states during O-to-ground transition in KR2, strongly suggest that Na^+^ release proceeds also on the path of least resistance without any notable structural rearrangements of the protein. This is supported by the fact that the retinal in the O-state is almost completely relaxed to its initial configuration. We also speculate that for the facilitation of the Na^+^ release (lowering of the energy barrier for the Na^+^ release against electrochemical gradient), the protein use the relay mechanism, in which the released ion replaces the ion bound at the oligomerization interface. The fact was also evidenced by the molecular dynamics simulations, described in the present manuscript. Current understanding of the Na^+^ in KR2 is summarized in the Supplementary Video 1.

## Materials and Methods

### Protein Expression and Purification

*E. coli* cells of strain SE1 (Staby™Codon T7, Eurogentec, Belgium) were transformed with the KR2 expression plasmid. Transformed cells were grown at 37° C in shaking baffled flasks in an auto-inducing medium ZYP-5052^28^ containing 100 mg/L ampicillin. When glucose level in the growing bacterial culture dropped below 10 mg/L, 10 µM all-*trans*-retinal (Sigma-Aldrich, Germany) was added, the incubation temperature was reduced to 20° C and incubation continued overnight. Collected cells were disrupted in M-110P Lab Homogenizer (Microfluidics, USA) at 25000 psi in a buffer containing 20 mM Tris-HCl pH 8.0, 5% glycerol, 0.5% Triton X-100 (Sigma-Aldrich, USA) and 50 mg/L DNase I (Sigma-Aldrich, USA). The membrane fraction of cell lysate was isolated by ultracentrifugation at 90000 g for 1 h at 4° C. The pellet was resuspended in a buffer containing 50 mM NaH_2_PO_4_/Na_2_HPO_4_ pH 8.0, 0.1 M NaCl and 1% DDM (Anatrace, Affymetrix, USA) and stirred overnight for solubilization. Insoluble fraction was removed by ultracentrifugation at 90000 g for 1 h at 4° C. The supernatant was loaded on Ni-NTA column (Qiagen, Germany) and KR2 was eluted in a buffer containing 50 mM NaH_2_PO_4_/Na_2_HPO_4_ pH 7.5, 0.1 M NaCl, 0.5 M imidazole and 0.1% DDM. The eluate was subjected to size-exclusion chromatography on 125 ml Superdex 200 PG column (GE Healthcare Life Sciences, USA) in a buffer containing 50 mM NaH_2_PO_4_/Na_2_HPO_4_ pH 7.5, 0.1 M NaCl, 0.05% DDM. Protein-containing fractions with the minimal A_280_/A_525_ absorbance ratio were pooled and concentrated to 60 mg/ml for crystallization.

### Measurements of pumping activity in E. coli cells

*E. coli* cells of strain C41(DE3) (Lucigen) were transformed with the KR2 expression plasmid. Transformed cells were grown at 37 °C in shaking baffled flasks in an autoinducing medium, ZYP-5052^28^ containing 100 mg/L ampicillin, and were induced at optical density OD600 of 0.7–0.9 with 1 mM isopropyl β-d-1-thiogalactopyranoside (IPTG) and 10 μM after induction, the cells were collected by centrifugation at 4,000g for 10 min and were washed three times with unbuffered salt solution (100 mM NaCl, and 10 mM MgCl_2_) with 30-min intervals between the washes to allow exchange of the ions inside the cells with the bulk. After that, the cells were resuspended in 100 mM NaCl solution and adjusted to an OD_600_ of 8.0. The measurements were performed on 3 ml of stirred cell suspension kept at 1 °C. The cells were illuminated for 5 min with a halogen lamp (Intralux 5000-1, VOLPI) and the light-induced pH changes were monitored with a pH meter (LAB 850, Schott Instruments). Measurements were of protonophore carbonyl cyanide 3-chlorophenylhydrazone (CCCP).

### Crystallization details and crystals preparation

The crystals were grown using the *in meso* approach^29, 30^, similarly to our previous work^26, 31, 32^. The solubilized protein in the crystallization buffer was added to the monooleoyl-formed lipidic phase (Nu-Chek Prep, USA). The best crystals were obtained using the protein concentration of 25 mg/ml. The crystals of monomeric (for D116N mutant) and pentameric (for WT protein and H30A mutant) forms were grown using the precipitate 1.0 M sodium malonate pH 4.6 and 1.2 M sodium malonate pH 8.0, respectively (Hampton Research, USA). Crystallization probes were set up using the NT8 robotic system (Formulatrix, USA). The crystals were grown at 22 °C and appeared in 2-4 weeks. Before harvesting, crystallization drop was opened and covered with 3.4 M sodium malonate solution, pH 8.0, to avoid dehydration. All crystals were harvested using micromounts (MiTeGen, USA) and were flash-cooled and stored in liquid nitrogen for further crystallographic analysis.

### Time-resolved visible absorption spectroscopy on KR2 crystals

The laser flash photolysis was performed similar to that described by Chizhov et al^33, 34^ with minor differences. The excitation system consisted of Nd:YAG laser Q-smart 450mJ with OPO Rainbow 420-680nm range (Quantel, France). For the experiments the wavelength of the laser was set 525 nm. Microcrystals of KR2 in the lipidic cubic phase were plastered on the 4x7mm cover glass. The thickness of the slurries was adjusted in order to give sufficient signal. The glass with crystal slurries was placed into 5x5mm quartz cuvette (Starna Scientific, China) filled with the buffer solution containing 3.4 M sodium malonate pH 8.0 and thermostabilized via sample holder qpod2e (Quantum Northwest, USA) and Huber Ministat 125 (Huber Kältemaschinenbau AG, Germany). The detection system beam emitted by 150W Xenon lamp (Hamamatsu, Japan) housed in LSH102 universal housing (LOT Quantum Design, Germany) passed through pair of Czerny–Turner monochromators MSH150 (LOT Quantum Design). The received monochromatic light was detected with PMT R12829 (Hamamatsu). The data recording subsystem represented by a pair of DSOX4022A oscilloscopes (Keysight, USA). The signal offset signal was measured by one of oscilloscopes and the PMT voltage adjusted by Agilent U2351A DAQ (Keysight).

### Spectroscopic characterization and accumulation of the intermediate state in KR2 crystals

Absorption spectra of KR2 in solution were collected using the UV-2401PC spectrometer (Shimadzu, Japan). The spectroscopic characterization of O-state build-up in KR2 crystals was performed at the icOS Lab located at the ESRF^35^. The same set up was established at the P14 beamline of the PETRAIII synchrotron source (Hamburg, Germany) for accumulation of the O-state in crystals for X-ray diffraction data collection. Also the same accumulation procedure was applied to crystals at icOS and P14 beamline. Briefly, UV-visible absorption spectra were measured using as a reference light that of a DH-200-BAL deuterium-halogen lamp (Ocean Optics, Dunedin, FL) connected to the incoming objective via a 200 µm diameter fiber, resulting in a 50 um focal spot on the sample, and a QE65 Pro spectrometer (Ocean Optics, Dunedin, FL) connected to the outgoing objective via a 400 µm diameter fiber. The actinic light comes from a 532 nm laser (CNI Laser, Changchun, P.R. China) coupled to a 1000 µm diameter fiber which is connected to the third objective whose optical axis is perpendicular to those of the ingoing and outgoing objectives. Ground states spectra (100 ms acquisition time averaged 20 times) were collected on crystals flash-cooled in liquid nitrogen and kept under a cold nitrogen stream at 100 K. In order to maximize the population of the O-state, a crystal was put under constant laser illumination at 100 K, the nitrogen stream was then blocked for 2 seconds, then the laser was switched off once the crystal is back at 100 K. For the accumulation of the O-state laser power density of 7.5 mW/cm^2^ at the position of the sample was used. The mean size of the crystals was 200x100x30 μm (Supplementary Fig 22). The plate-like crystals were oriented so that the largest plane (200x100 μm to the size of 500x500 ^2^μ) was as perpendicular to the laser beam. The laser beam was focused to the size of 500x500 μm^2^ (1/e^2^). A UV-visible absorption spectrum was then recorded to show the red-shifted absorption maximum characteristic of the O-state of KR2. The crystals with accumulated intermediate state were then stored in liquid nitrogen and transported to the PETRAIII, Hamburg, Germany for the X-ray experiments and showed the same structure as that obtained using crystals with the O-state, accumulated directly at the P14 beamline of PETRAIII.

### Acquisition and treatment of diffraction data

X-ray diffraction data of D116N and H30A mutants were collected at the beamlines ID23-1 and ID29 of the ESRF, Grenoble, France, using a PILATUS 6M and EIGER 16M detectors, respectively. X-ray diffraction data of the KR2 O-state at 100 and 293K (room temperature, RT) was collected at the P14 beamline of the PETRAIII, Hamburg, Germany, using EIGER 16M detector. For the collection of the X-ray diffraction data at RT the crystals in the cryoloop were placed on the goniometer of P14 beamline and maintained in the stream of the humid air (85% humidity). The stream of humid air was provided by the HC humidity controller (ARINAX, France). For activation of the proteins in crystals and obtaining the structure of the O-state at RT the laser flash was synchronized with the X-ray detector. The laser was illuminating the crystal only during X-ray data collection to avoid drying and bleaching. The crystals were rotated during the data collection and laser illumination. Diffraction images were processed using XDS^36^. The reflection intensities of the monomeric form of D116N mutant were scaled using the AIMLESS software from the CCP4 program suite^37^. The reflection intensities of all the pentameric forms were scaled using the Staraniso server^38^. There is no possibility of twinning for the crystals. For the both structures of KR2 at RT diffraction data from three crystals was used (Table S1). In all other cases, diffraction data from one crystal was used. The data statistics are presented in the Table S1.

### Structure determination and refinement

Initial phases for the pentameric structures were successfully obtained in the C222_1_ space group by molecular replacement (MR) using MOLREP^39^ using the 6REW structure as a search model. Initial phases for monomeric KR2-D116N were successfully obtained in the I222 space group by MR using the 4XTL structure as a search model. The initial MR models were iteratively refined using REFMAC5^40^, PHENIX^41^ and Coot^42^.

### Molecular dynamics simulations

The simulation system consisted of a KR2 pentamer in the O-state with sodium ions in SBC2 and cocrystallized water molecules. The proteins was embedded in a POPC bilayer (256 lipids) and then solvated with TIP3P water with a Na+/Cl− concentration of 150 mM using the CHARMM-GUI web-service^43^. The simulation box contained 94817 atoms in total. All ionizable amino acids were modeled in their standard ionization state at pH 8, including D116 and D251 which were modeled charged.

The CHARMM-GUI recommended protocols were followed for the initial energy minimization and equilibration of the system. The atoms of protein and lipids in the system were subjected to a harmonic positional restraint and 5000 steps of steepest descent minimization followed by two 25 ps equilibration steps in the NVT ensemble using Berendsen thermostat and one 25 ps and three 50 ps equilibration steps in the NPT ensemble using Berendsen thermostat and barostat. During all equilibration steps, the force constants of the harmonic positional restraints were gradually reduced to zero. The system was further equilibrated for 10 ns in the NPT ensemble with Nose-Hoover thermostat and Parrinello−Rahman barostat, which were also used for the further production simulations. The temperature and pressure were set to 303.3 K and 1 bar with temperature and pressure coupling time constants τt = 1.0 ps−1 and τp = 0.5 ps−1, respectively.

All MD simulations were performed with GROMACS version 2018.1^44^. The time step of 2 fs was used for all the simulations except for the early steps of equilibration. The CHARMM36 force field^45^ was used for the protein, lipids, and ions. Parameters for retinal bound to lysine were adapted from^46^.

In order to investigate the putative sodium translocation pathways, metadynamics approach (metaMD) was employed^47^. This method is based on biasing of the potential surface via addition of repulsive functions (“hills”, typically Gaussians) which force the investigated molecular system to explore its configurational space broader and faster than in a regular unbiased MD simulation. We used the PLUMED plugin for GROMACS to perform metaMD simulations^48^. The projections of the vector connecting the sodium ion and its original position onto x, y and z directions were used as the collective variables (i.e., 3 CVs were used). In order to prevent sodium passage back to the cytoplasmic side we applied a flat-bottom potential (k=1000 kJ/mol/nm^2^) in the normal to the membrane direction, which discouraged ion moving towards the cytoplasmic side. Also, harmonic restrains were applied to all protein C atoms above the C atom of K255 (in the direction of CP) to prevent the overall motion of the protein complex. The deposition rate for hills was 0.5 ps; the width and height of deposited hills were equal to 0.05 nm and 1 kJ/mol, respectively. The simulations were continued until the exit of ion from the protein interior was observed (typically, during 5-20 ns). We have carried out 10 metaMD runs in total, 2 replicates for each of the 5 protomers of the KR2 pentamer.

## Acknowledgments

We acknowledge the Structural Biology Group of the European Synchrotron Radiation Facility (ESRF) and The European Molecular Biology Laboratory (EMBL) unit in Hamburg at Deutsche Elektronen-Synchrotron (DESY) for granting access to the synchrotron beamlines

## Author contributions

CB, SV and DZ expressed and purified the proteins; TB supervised the expression and purification; RA and KK crystallized the proteins; DZ helped with crystallization; KK collected absorption spectra from crystals and performed cryo-trapping of the intermediate; AR and PC supervised the absorption spectra collection; DV performed flash photolysis experiments on KR2 crystals and processed the data; KK helped with the flash photolysis experiments; KK collected the diffraction data with the help of RA, DZ and solved the structures; EZ and EM processed the serial crystallography data; IG supervised structure refinement and analysis; GB and AP helped with data collection; PO performed the molecular dynamics simulations; IS prepared the systems for simulations; MR performed oligomerization analysis of the proteins; AA, VB, EB and GB helped with data analysis, VG supervised the project; KK and VG analyzed the results and prepared the manuscript with input from all the other authors.

## Supplementary Information

### Supplementary text

#### Key role of S-N-D triad in ion translocation

The S70-N112-D116 (S-N-D) triad, which is completely conserved within NaRs and comprises the core of the Na^+^ binding site in the middle part of the protein, is very similar in respect to the residues composition and their relative location to the central gates (CGs) of channelrhodopsins (Supplementary Fig 15). Indeed, the CGs of the native cation channelrhodopsin-2 (*Cr*ChR2)^50^, chimeric cation channelrhodopsin (C1C2)^51^ and also natural anion channelrhodopsin (*Gt*ACR1)^52, 53^ are composed of the S63-N258-E90, S102-N297-E129 and S43-N239-E68 triads (Supplementary Fig 15). Although the residues are located in the different helices in comparison to the KR2, they form similar overall conformation. CGs serve as constriction sites in the central part of the channels, and are important for the ion selectivity^54^. This allows us to suggest that the transient Na^+^ binding site is important for KR2 selectivity. It cannot be also excluded that CGs may act as transient binding sites in channelrhodopsins during ion translocation.

Moreover, in 2015 the evolutional relationship of microbial light-driven Na^+^ pumps and class A G protein-coupled receptors (GPCRs) was studied, based on the existing structures of the ground state of KR2^55^. Presented here structure of the O-state with Na^+^ bound inside the protein supports the high similarity of the Na^+^ binding sites in KR2 and GPCRs. In particular, in both cases the sites are located near helices B and C (TM2 and TM3 in GPCRs), and formed by the similar to the S-N-D triad of KR2 set of residues (D95, N131 and S135 in case of human δ oid receptor) (Supplementary Fig 16). This opens the way for more accurate analysis of the interconnection between two highly important families.

Last but not least, the Na^+^ binding site inside KR2 is also very similar to that of another type of Na^+^-transporting proteins - Na^+^ ATP synthases. For instance, it is almost identical to that of the c11-ring of the Na^+^ ATP synthase from *Ilyobacter tartaricus* (Supplementary Fig 16).

#### The second Na^+^ identified at the KR2 surface in the ground state

Recently we solved crystal structure of KR2 pentameric Na^+^-pumping state under physiological conditions at 2.2 Å resolution^4^. In the frame of present work, we improved the resolution of the model to 2 Å. Therefore, we present here more complete structure of the KR2 resting state at 2 Å (Supplementary Table 1).

While the inner region of the protein protomers is the same as previously reported, we identified numerous of water molecules at the KR2 surface. Moreover, we observe a second Na^+^ binding site at the pentamer surface (Supplementary Fig 17). The additional Na^+^ is coordinated by the main chain oxygen of S100 residue and 5 water molecules with the mean Na-O distance of 2.5 Å (Supplementary Fig 17). Notably, the Na^+^ is located close to the putative ion release cavity 2 (pIRC2). As the Na^+^ was not identified at lower resolution, and its B-factor is considerably higher than that of the previously reported Na^+^ (52 and 21 Å, respectively), we suggest that this ion may play a key role in the relay mechanism of Na^+^ release from the pIRC2 to the extracellular space (Supplementary Fig 17). Indeed, the loosely bound Na^+^ may be released to the bulk from the surface upon O-to-ground transition and substituted by the ion transported in the current cycle from the pIRC2.

#### Crystal structure of the ground and the O-states of KR2 at room temperature

In order to check that freezing does not affect the KR2 conformations in the ground and the O-state we collected diffraction data on KR2 at 293 K using single crystals placed in the stream of humid air (relative humidity of 85%) and solved the structures of the dark (ground) and illuminated (49% ground : 51% O-state) at 2.5 and 2.6 Å, respectively. To accumulate the O-state in crystals, we illuminated them continuously by 532 nm laser during X-ray data collection. This approach allows accumulation of the dominant intermediate of protein photocycle, which in the case of KR2 is the O-state. Analysis of the electron density maps and occupancies refinement indicated that such procedure results in the 51% occupancy of the O-state in crystals.

Overall, the structures of the ground and the O-states of KR2 at 100 and 293 K are nearly identical (Supplementary Fig 18, 19). The RMSD between the structures of the ground state of KR2 at 100 and 293K is 0.2 Å, and between those of the O-state it is also 0.2 Å. Comparison of the KR2 structures identified slight shifts of the positions of the E-F and F-G loops and also cytoplasmic parts of helices E and F by only 0.4 Å (Supplementary Fig 18). The orientations of all key residues and the arrangement of water molecules inside the protein are similar in models at 100 and 293K (Supplementary Fig 19).

#### Double conformation of the RSBH^+^ in D116N mutant

The electron densities around retinal and K255 in the D116N mutant suggest the coexistence of two alternative orientations of the RSBH^+^ (Supplementary Fig 20). In one of them, similar to the O-state and ‘compact’ conformation of KR2, RSBH^+^ is pointed towards N116. However, there is no hydrogen bond between them and the distance between RSBH^+^ and N116 is 4.3 Å. Likely, there is a hydrogen bond to N112 in this conformation. In the second conformation, RSBH^+^ is shifted closer to D251 (3.1 Å) and may form a hydrogen bond with the residue (Supplementary Fig 6D, 21). In both conformations, retinal remains in the all-*trans* configuration (Supplementary Fig 20). Importantly, such organization of the RSBH^+^ region is in line with the existing spectroscopic and nuclear magnetic resonance (NMR) data on D116N mutant. Indeed, it was shown, that the hydrogen bond between the RSBH^+^ and D116^-^ exists in only a fraction of this KR2 variant. Moreover, as it was demonstrated by the NMR studies, the RSBH^+^ is in a multiple conformations in D116N^56^. Therefore, we refined our crystallographic data on the mutant with the both alternative RSBH^+^ orientations in the final model (Supplementary Fig 20).

#### The basis of the KR2 pentamer dissociation at acidic pH

We showed previously that the oligomeric state of the KR2 is pH-dependent in the detergent micelles and also in the crystals grown from the lipidic cubic phase^3, 4^. KR2 is organized into pentamers at pH values higher than 6-6.5, while the monomers are found at acidic pH. The basis of pentamer disruption at low pH remains unclear. One of the hypotheses is the influence of pH on the rechargeable residues of the oligomerization interface, such as H30. It was demonstrated that pentameric assembly is disturbed in H30K and H30L mutants^4^. However, the H30A mutant remained pentameric^12^. This suggests another mechanism of oligomer dissociation. As it is shown, with the pH decrease not only the pentameric assembly is affected, but also protonation of D116 occurs. For the wild type protein, the shift from the pentameric to monomeric state appears detergent at pH 5-6, which is also close to the pKa of D116. Consequently, it is natural to suggest that the protonation of D116 may influence the oligomeric state of the protein and lead to the pentamer dissociation. To check the hypothesis, we studied oligomerization of the D116N in detergent micelles using size-exclusion chromatography (SEC). The mutant D116N imitates the protein with protonated D116 at all pH values. We showed that at pH 8.0, where KR2 forms pentamers, D116N is observed in several oligomeric states. Notable portions of both smaller and bigger oligomers are present in the solution. As follows from the analysis of the D116N oligomerization dependence on pH, the smaller oligomers dominate at pH 6, but some intermediate-sized oligomers (smaller than pentamers of the wild type protein) appear as pH is lowered to 4.3 (Supplementary Fig 21). Altogether, this allows us to suggest that the protonation of D116 and presumably binding of Na^+^ near D116 in the O-state affects the oligomeric state of KR2 and is one of the driving forces of the destabilization of the KR2 pentamer. Consequently, this additionally supports the fact that KR2 protomers are distorted in the O-state and pentamerization interface could also be affected. Thus, pentameric assembly not only plays a key role in the organization of the ‘expanded’ conformation, important for the Na^+^ release, but also stabilizes the proteins in the O-state with Na^+^ bound in close proximity of the RSBH^+^.

**Supplementary Figure 1.**
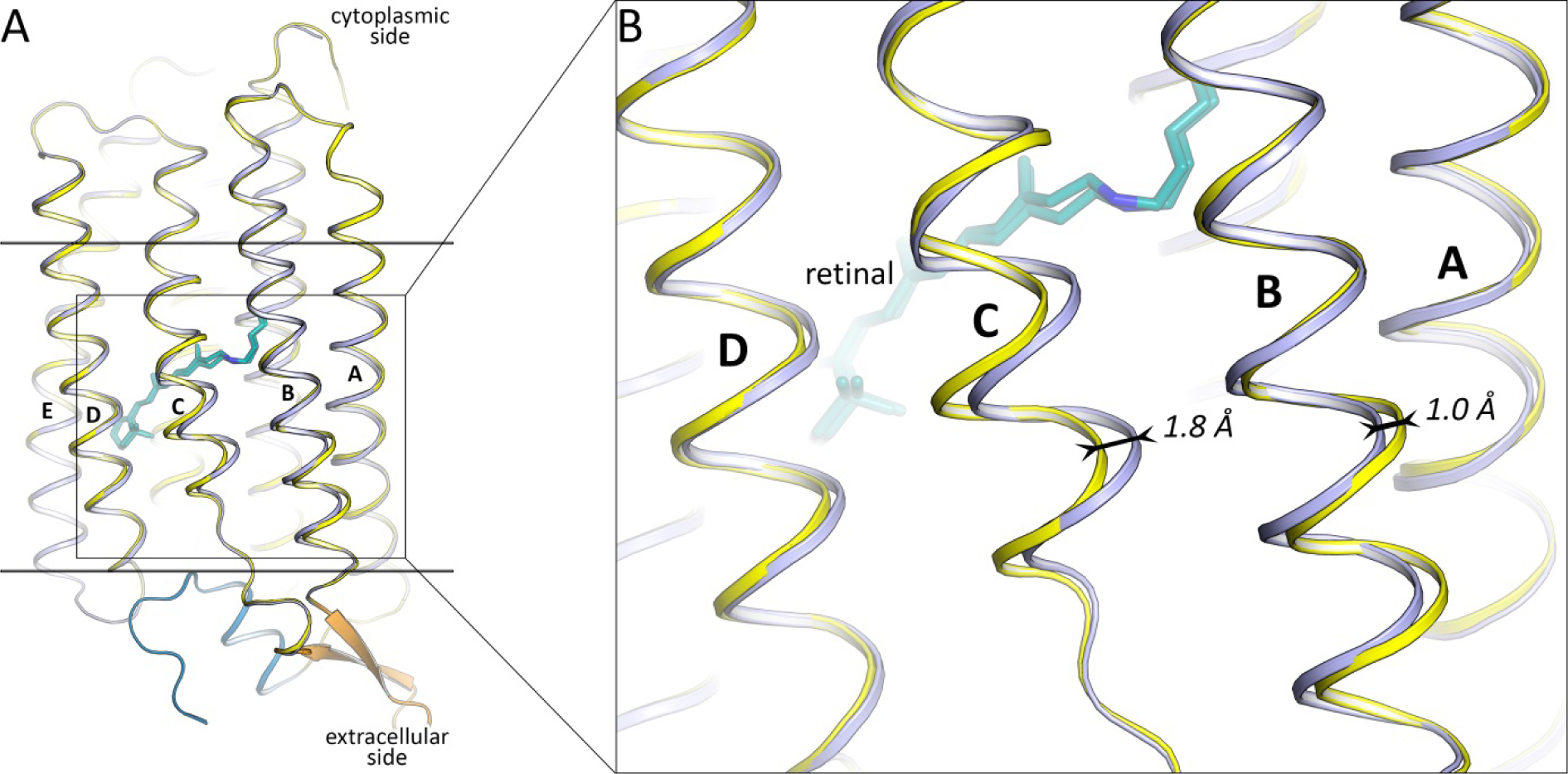
Structural alignment of KR2 protomers in the ground (yellow) and the O-(blue) states. **A.** Overall alignment. N-terminal α helix and N terminus are colored blue. BC loop, containing the β sheet, is colored orange. **B.** Enlarged view of the most notable rearrangements in protomer backbone. Retinal cofactor is colored teal. Membrane core boundaries are shown with black lines. Helices are indicated with capital letters. The shifts of extracellular parts of helices B and C are demonstrated with black arrows.

**Supplementary Figure 2.**
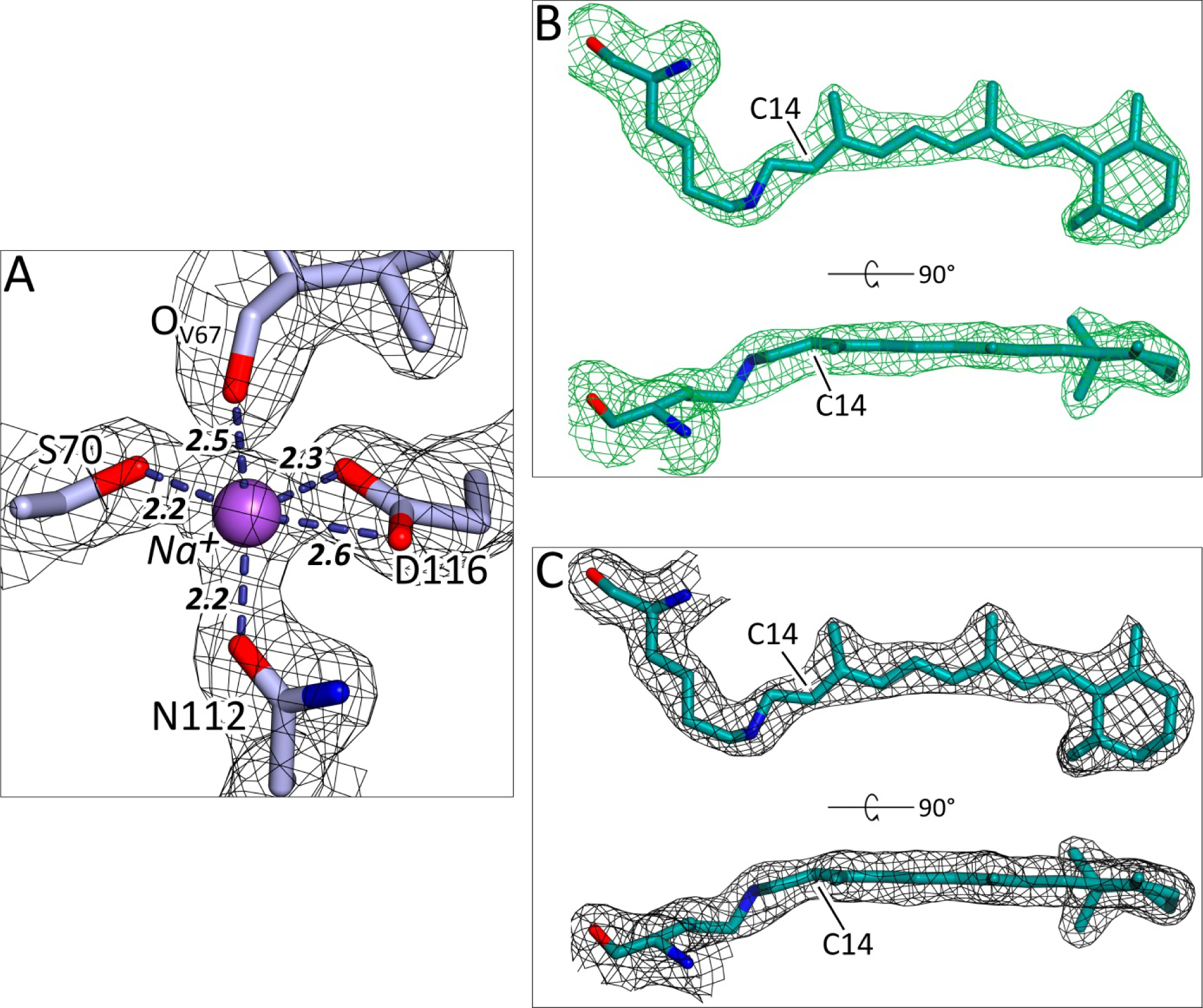
Examples of electron densities of the KR2 O-state. **A.** 2F_o_-F_c_ electron density maps around sodium ion binding site contoured at the level of 1.5 σ Sodium ion is shown with violet sphere. Hydrogen bonds, coordinating the sodium ion are shown with blue dashed lines. **B.** Polder^20^ maps for K255 and retinal cofactors of all five protomers of KR2 O-state structure. Maps are contoured at the level of 4.0 σ K255 and retinal cofactor contoured at the level of 1.5 σ **C.** 2F_o_-F_c_ electron density maps around Retinal and K255 are colored teal. C14 atoms of retinal are indicated. The lengths of the hydrogen bonds are shown with bold italic numbers and are in Å.

**Supplementary Figure 3.**
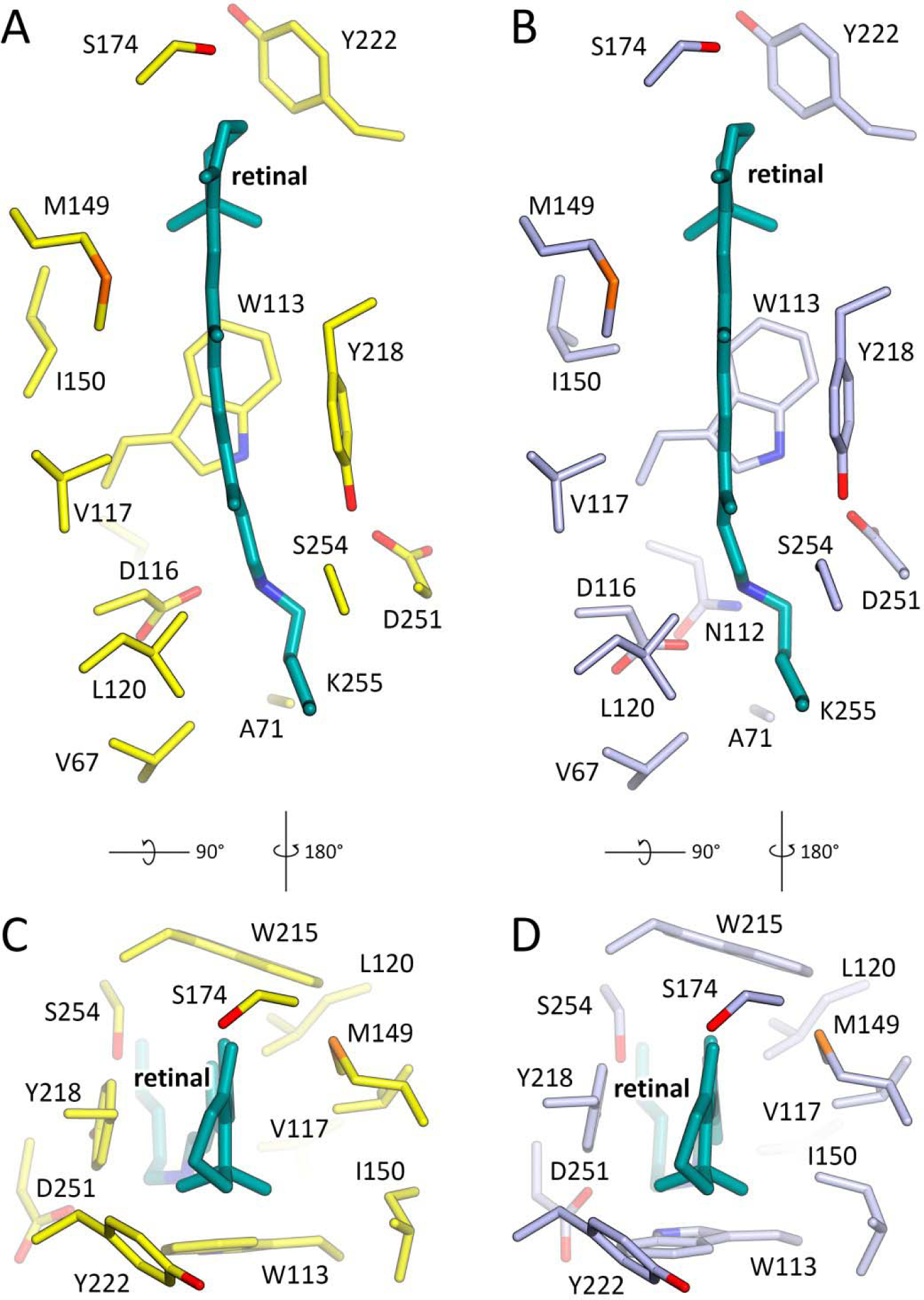
Retinal binding pockets of ground (yellow) and O (blue) states. **A, B.** View from the cytoplasmic side. **C, D.** View from the side of β onone ring of the retinal molecule. Retinal cofactor is colored teal.

**Supplementary Figure 4.**
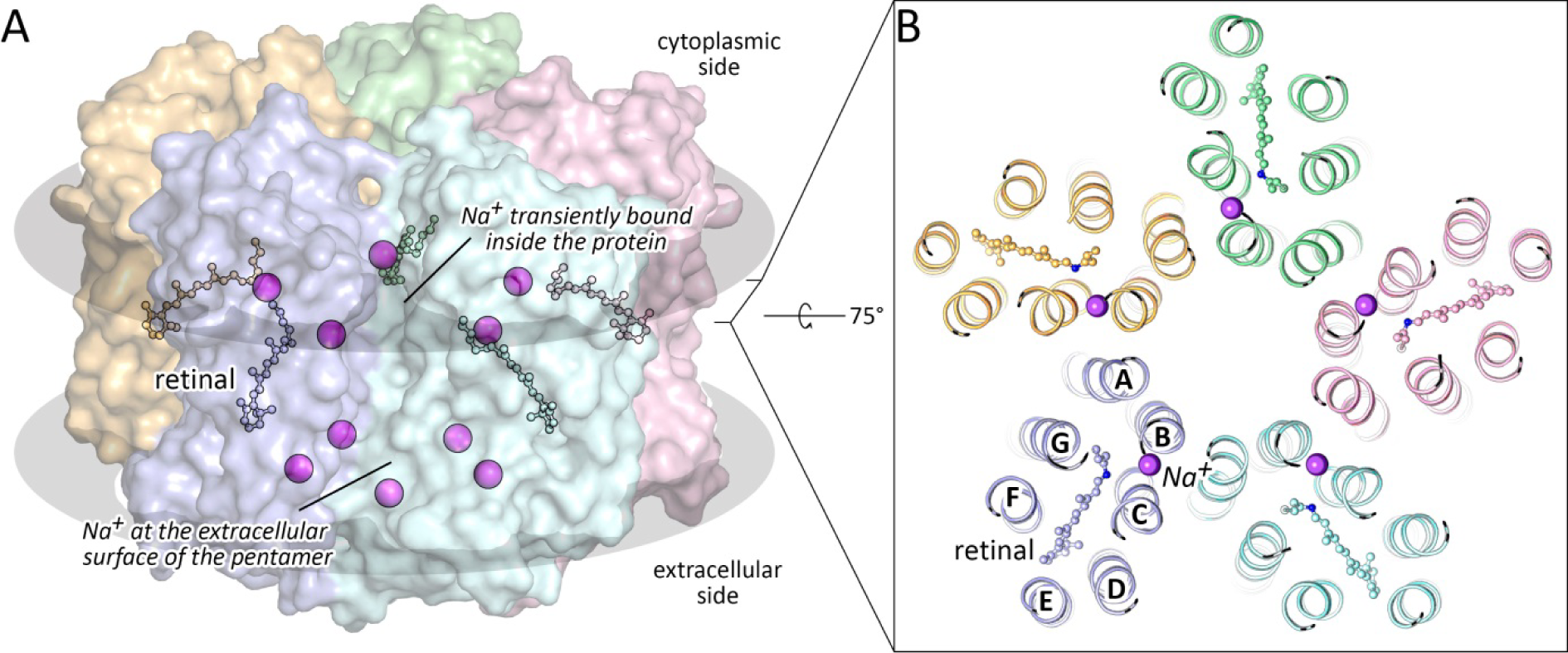
KR2 pentamer in the O-state and binding site of Na^+^. **A.** Side view of the KR2 pentamer, shown in surface representation. Membrane core boundaries are shown with black ellipses. **B.** Section view from the cytoplasmic side on the central part of the KR2 pentamer. Sodium ion is placed between helices B and C. Helices are indicated with capital letters. Sodium ions are colored violet. Retinal molecules and covalently bound Lys255 side chains are shown in balls and sticks representation.

**Supplementary Figure 5.**
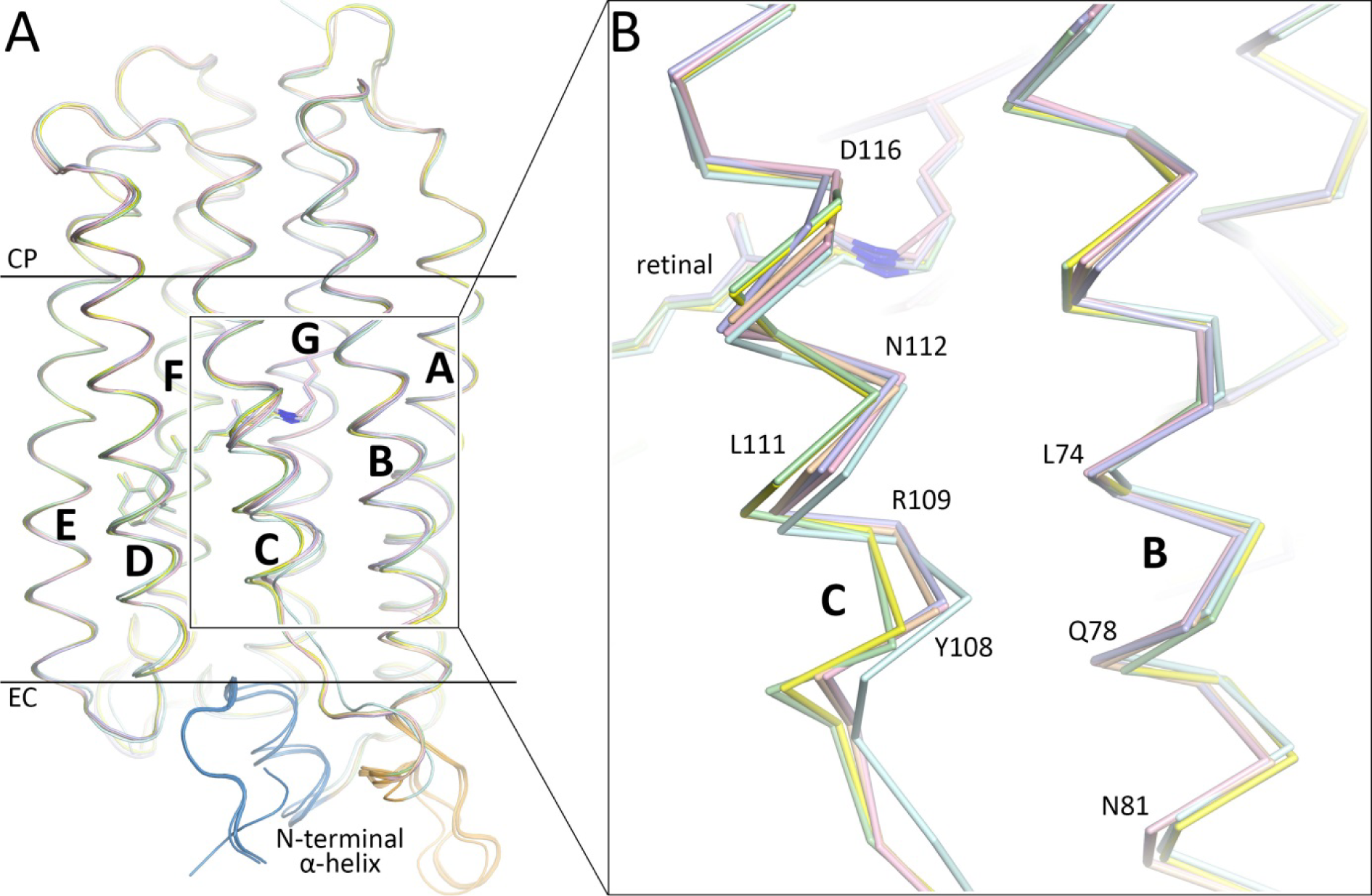
Structural alignment of KR2 protomers in different states. A. Overall alignment. N-terminal α helix and N terminus are colored blue. BC loop, containing the β-sheet, is colored orange. Membrane core boundaries are shown with black lines. **B.** Enlarged β view of the most notable rearrangements in protomer backbone. Helices are indicated with capital bold letters. Ground state of KR2 (the ‘expanded’ conformation, PDB ID: 6REW) is colored yellow. O-state of KR2 (present work) is colored light-blue. KR2-D116N (present work) is colored pink. KR2-H30A (present work) is colored green. Ground state of KR2 (the ‘compact’ conformation, PDB ID: 4XTN, chain ‘I’) is colored orange. Ground state of monomeric KR2 (PDB ID: 4XTL) is colored cyan.

**Supplementary Figure 6.**
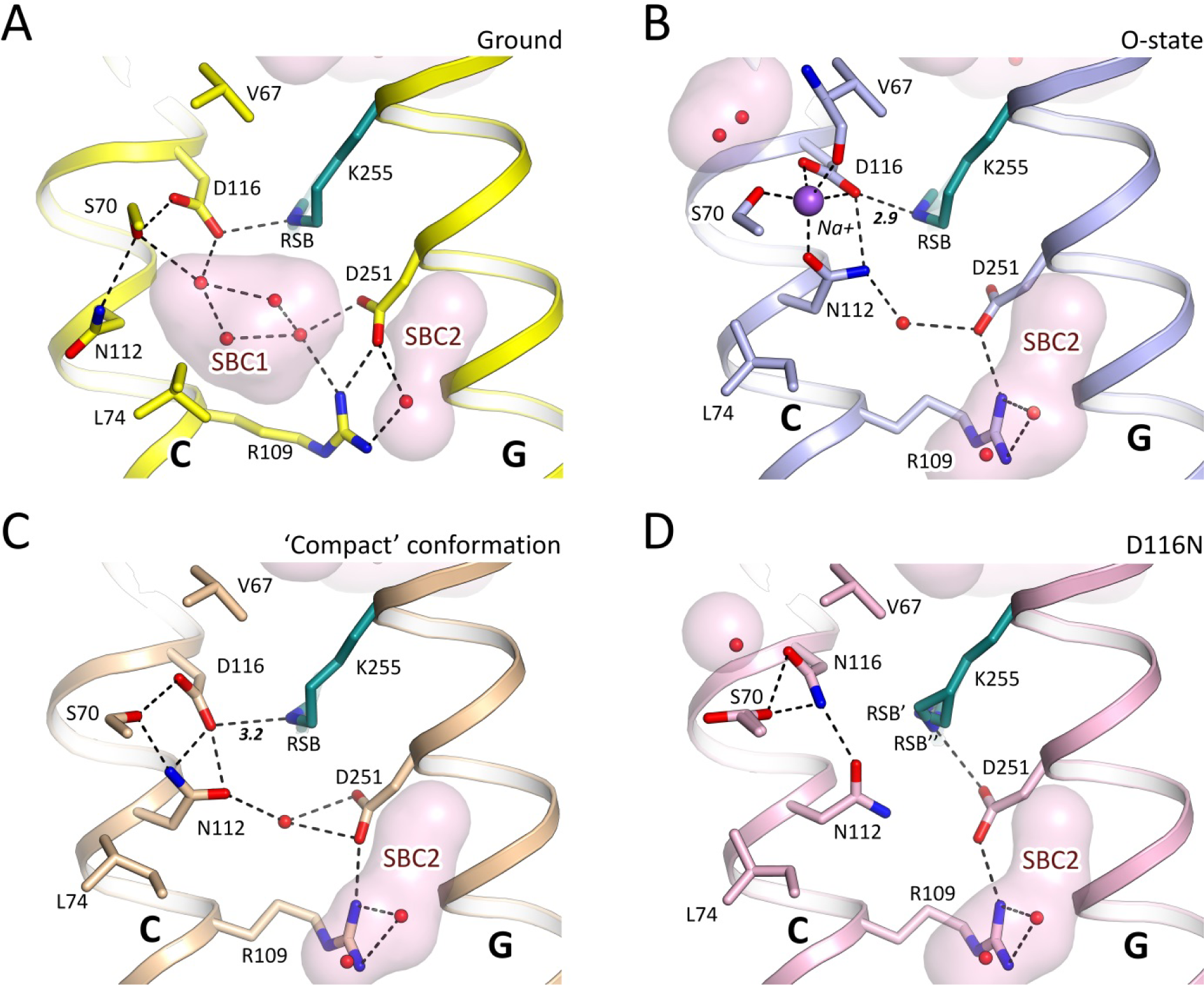
RSB region of the KR2 in different states. **A.** Ground state of KR2 (PDB ID: 6REW) **B.** O-state of KR2 (present work). **C.** The ‘compact’ conformation of KR2 (PDB ID: 4XTN, chain ‘I’). **D.** D116N mutant of KR2 (present work). Cavities are calculated using HOLLOW^49^ and shown with pink surfaces. Retinal cofactor is colored teal. Water molecules are shown with red spheres. Na^+^ is shown with a purple sphere. Hydrogen bonds are shown with black dashed lines. The lengths of the D116-RSB hydrogen bond are shown with bold italic numbers and are in Å. Helices C and G are indicated with capital bold letters. Helices A and B are hidden for clarity.

**Supplementary Figure 7.**
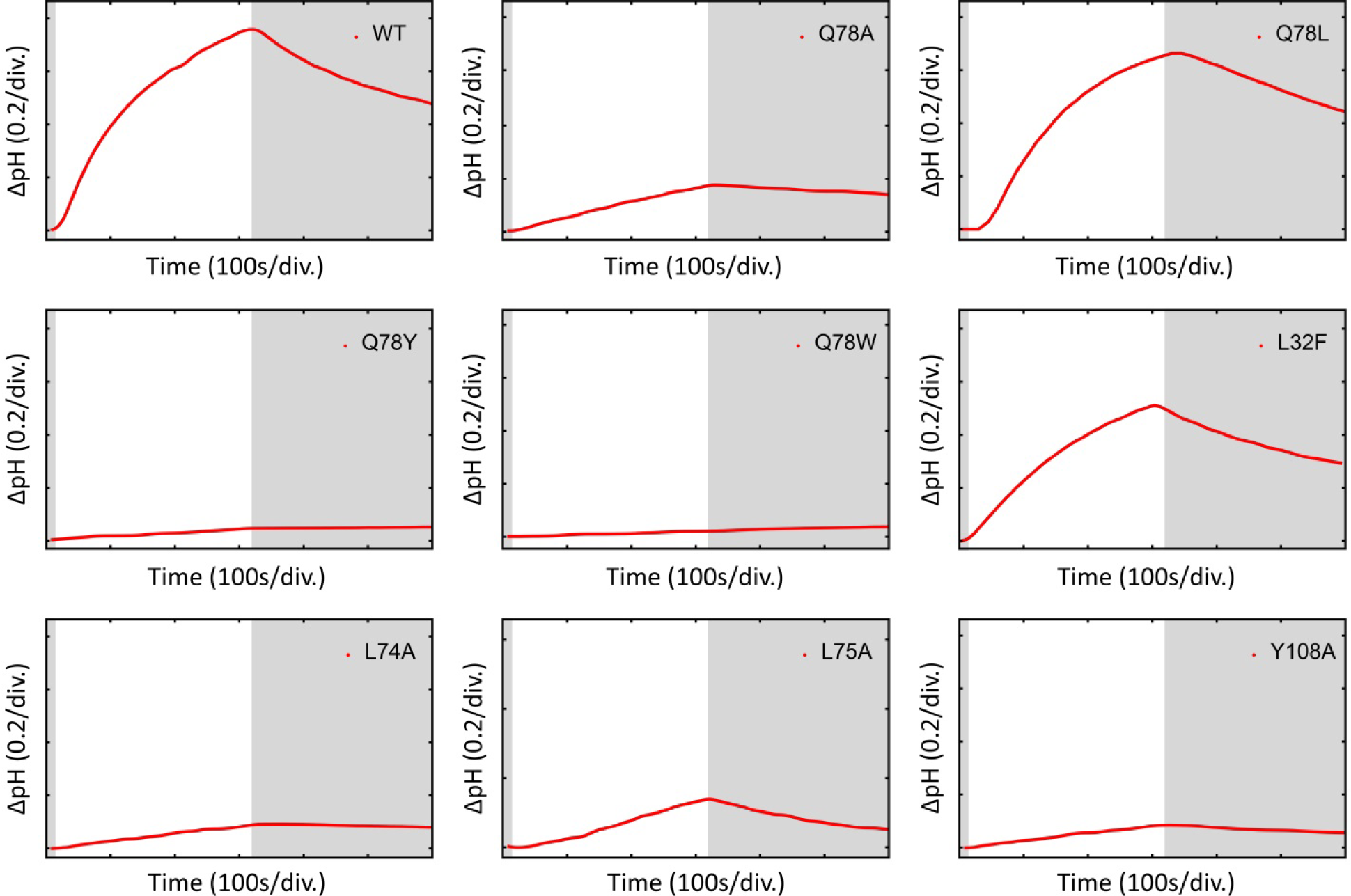
*E. coli* activity tests of KR2 and its mutants. pH changes upon illumination in the media containing KR2-expressing *E. coli* cells. The solutions contain 100 mM NaCl and 30 µM CCCP (magenta). The cells were illuminated for 300 s (light area on the plots).

**Supplementary Figure 8.**
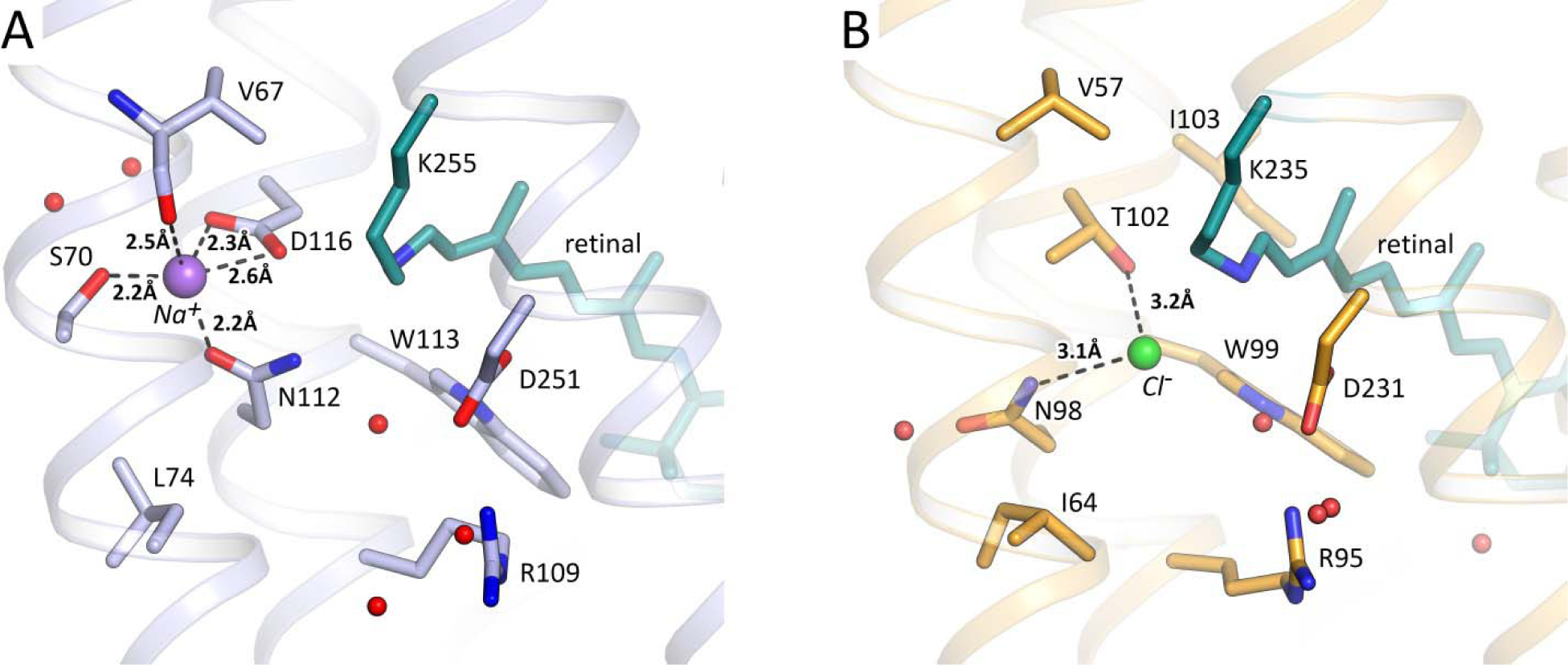
Na^+^ and Cl^-^ binding sites of the ion-pumping bacterial rhodopsins. **A.** O-state of KR2 (present work). **B.** Ground state of the chloride-pumping *Nonlabens marinus* S1-08 rhodopsin (PDB ID: 5ZTK^24^). Retinal cofactor is colored teal. Water molecules are shown with red spheres. Na^+^ is shown with a purple sphere. Cl^-^ is shown with a green sphere. Distances are shown with black dashed lines. The length of the D116-RSB hydrogen bond are shown with bold italic numbers and are in Å. Helix A is hidden for clarity.

**Supplementary Figure 9.**
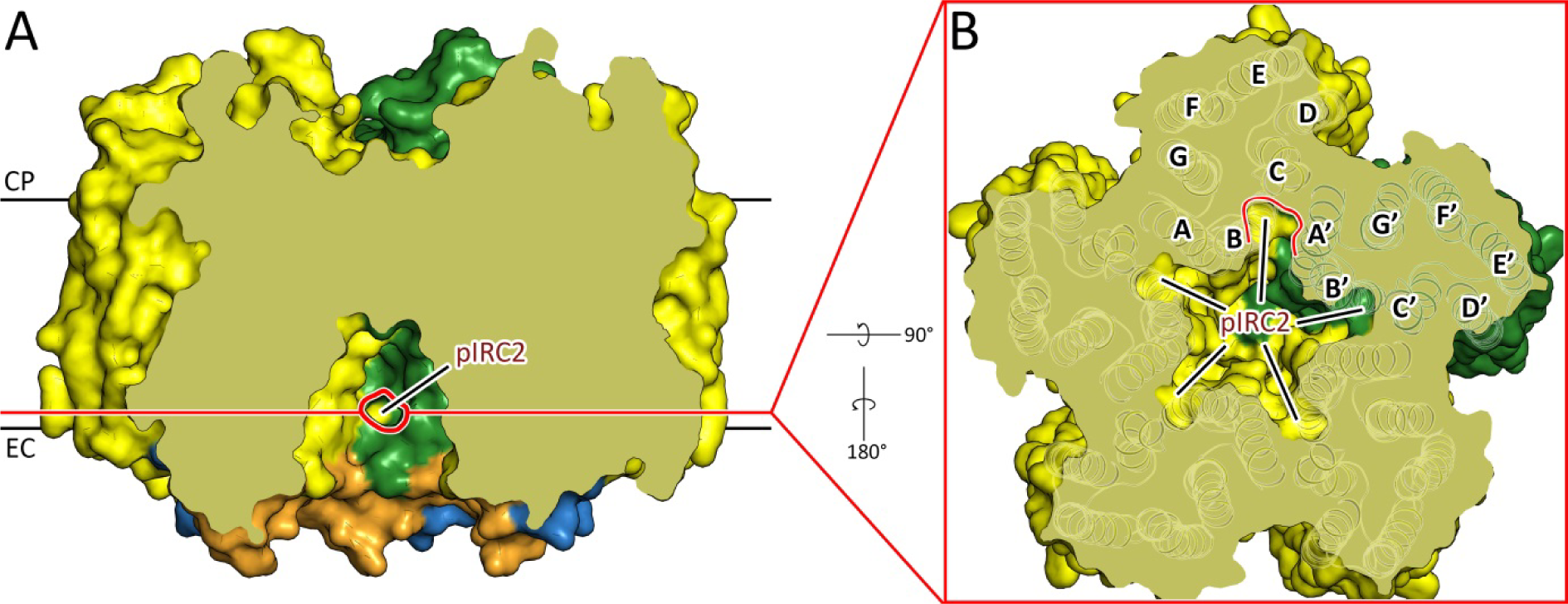
Putative ion-release cavity 2 (pIRC2) of the ground state of KR2. **A.** Side section view of KR2 pentamer. Red ellipse-like shape contours the pore in the protein surface leading to the pIRC2 from the concave aqueous basin formed in the central pore of KR2 pentamer. Membrane core boundaries are shown with black lines. **B.** Section view from the extracellular side at the level of the pIRC2. pIRC2 of each protomer is formed by helices B, C and BC loop and A’ from adjacent protomer. The pIRC2 from the section A is also contoured with red line. Helices are signed with capital bold letters.

**Supplementary Figure 10.**
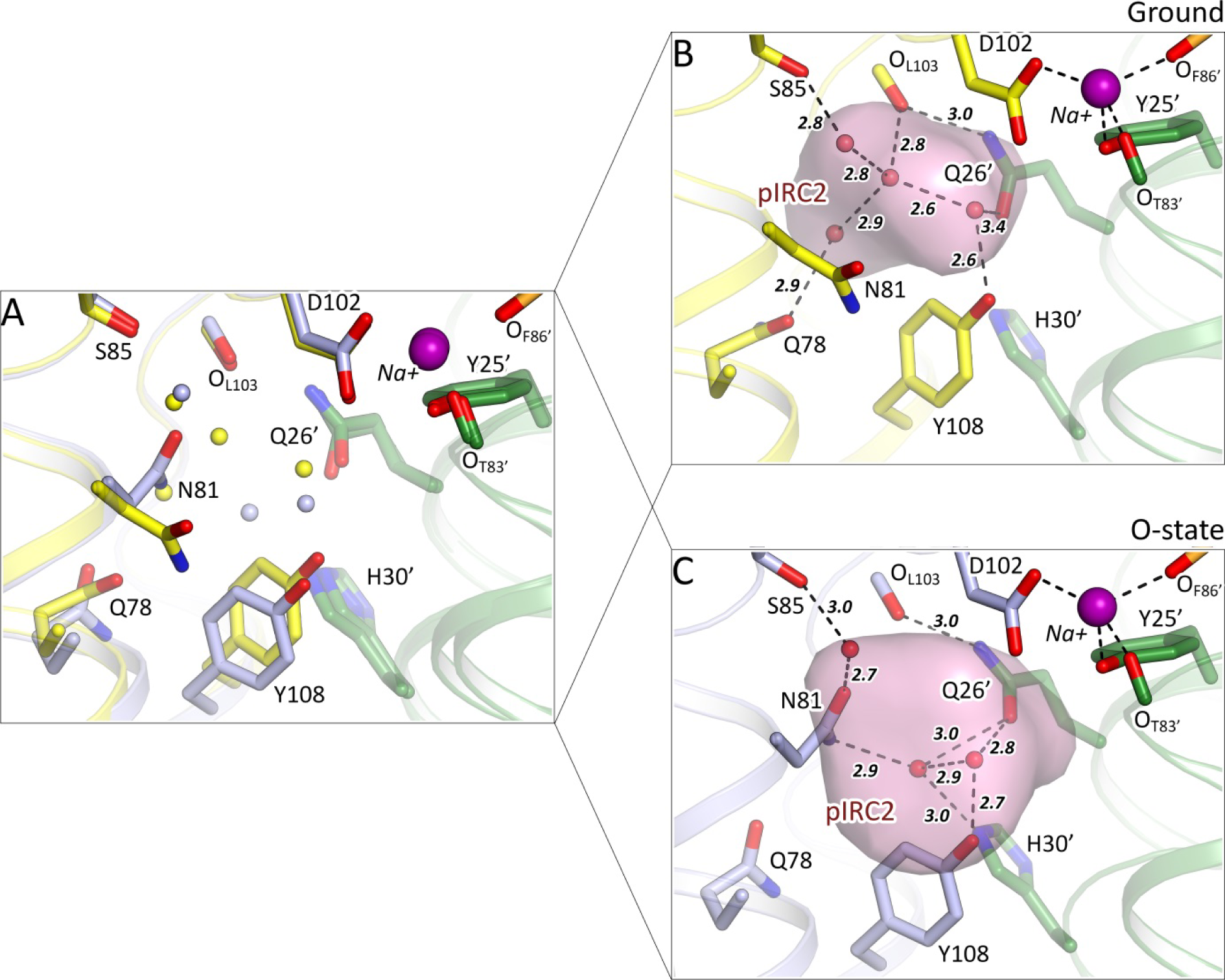
Putative ion-release cavity 2 (pIRC2) of the ground and O-states of KR2. **A.** Structural alignment of the ground (yellow, PDB ID: 6REW) and the O- (blue, present work) states. **B.** Detail view of the pIRC2 in the ground state. **C.** Detail view of the pIRC2 in the O-state. Na^+^ at the extracellular surface of KR2 pentamer are shown with purple spheres. Adjacent protomer is colored green. Hydrogen bonds are shown with black dashed lines. The length of the hydrogen bonds are shown with bold italic numbers and are in Å. Water molecules are shown with small spheres and colored yellow and blue in section A and red in sections B and C.

**Supplementary Figure 11.**
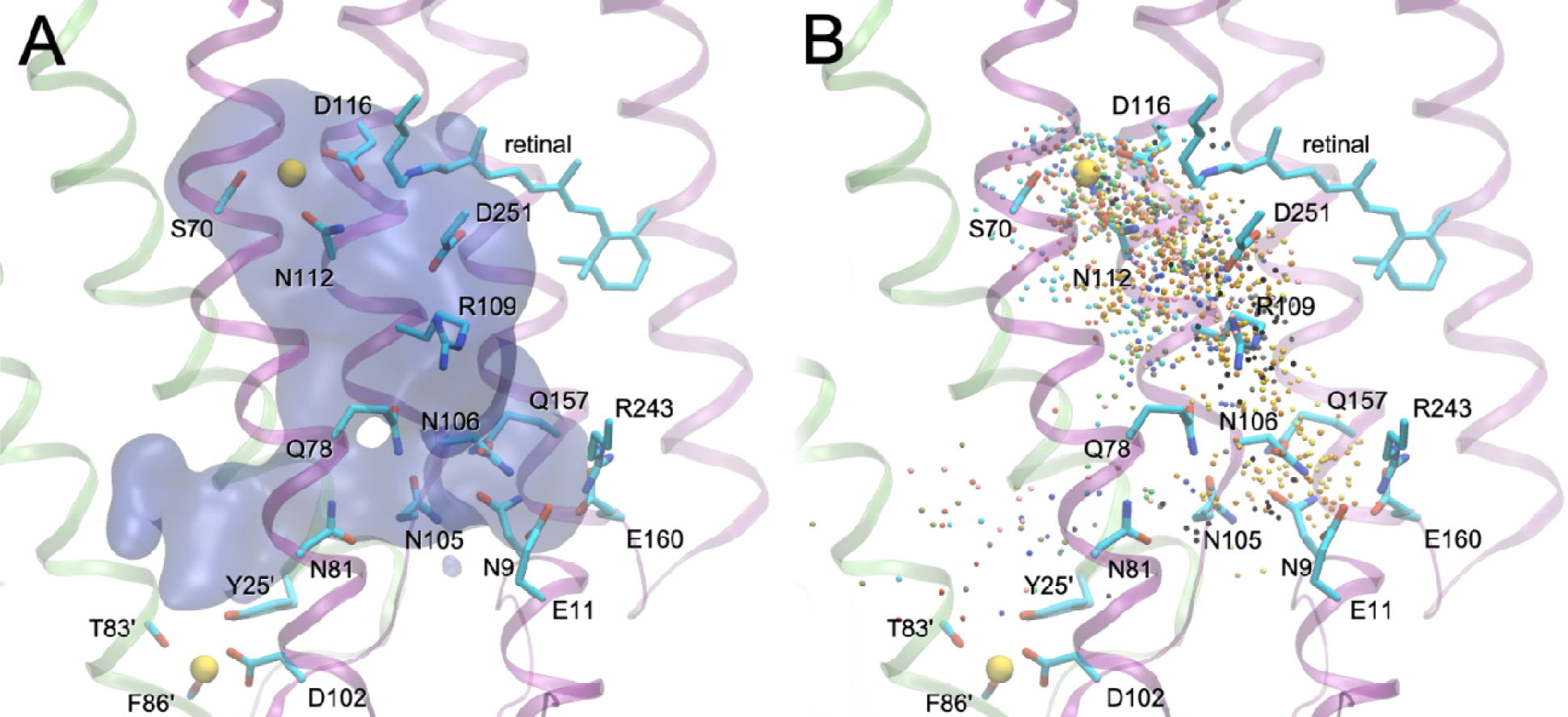
Simulated trajectories of Na^+^ release identified by molecular dynamics simulations. **A.** Density surface corresponding to the volume accessible to Na^+^. **B.** Positions of sodium taken every 100 ps. Each trajectory is shown in a different color; trajectories where sodium exits via pIRC1 are shown in yellow and orange. Two protomers of KR2 pentamer are rendered as purple and green ribbons. X-ray orientations of key amino acid sidechains are shown.

**Supplementary Figure 12.**
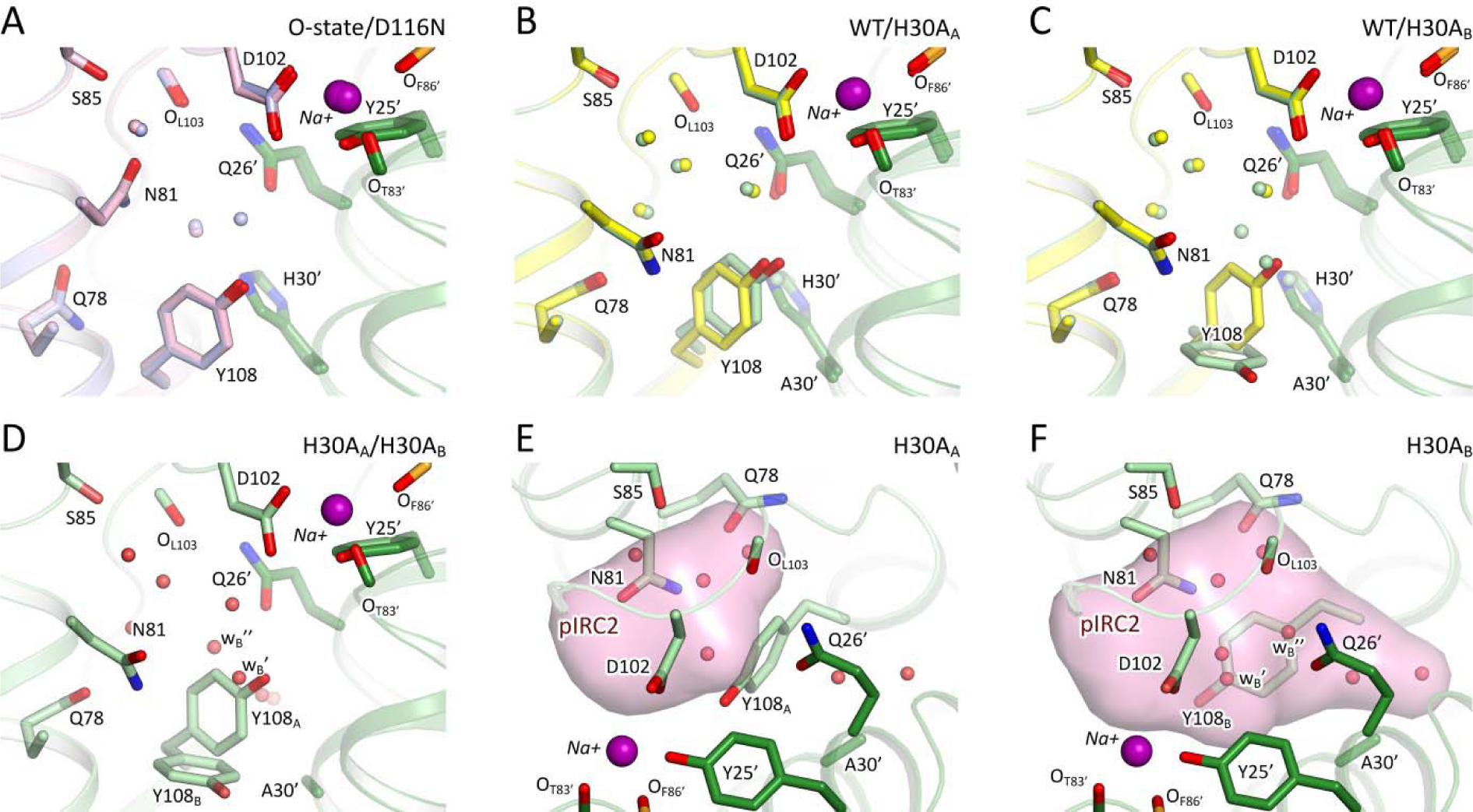
Putative ion-release cavity 2 (pIRC2) of the D116N and H30A mutants of KR2. **A.** Structural alignment of the O-state of the WT KR2 (blue, present work) and the D116N ground state (pink, present work). The conformations are nearly identical. **B.** Structural alignment of the ground states of the WT KR2 (yellow, PDB ID: 6REW) and conformation A of H30A mutant (H30A_A_, light green, present work). **C.** Structural alignment of the ground states of the WT KR2 (yellow, PDB ID: 6REW) and conformation B of H30A mutant (H30A_B_, light green, present work). **D.** Structural alignment of the ground states of the conformations A and B of H30A mutant. Additional water molecules, appearing in the structure of the H30A_B_ are identified as w_B_’ and w_B_’’. **E.** Detail view of the pIRC2 of the H30A_A_. **F.** Detail view of the pIRC2 of the H30A_B_. pIRC2 is notably enlarged in the H30A_B_. Na^+^ at the extracellular surface of KR2 pentamer are shown with purple spheres. Adjacent protomer is colored dark green. Water molecules are shown with small spheres and colored blue and pink in section A, yellow and light green in sections B and C and red in sections D-F.

**Supplementary Figure 13.**
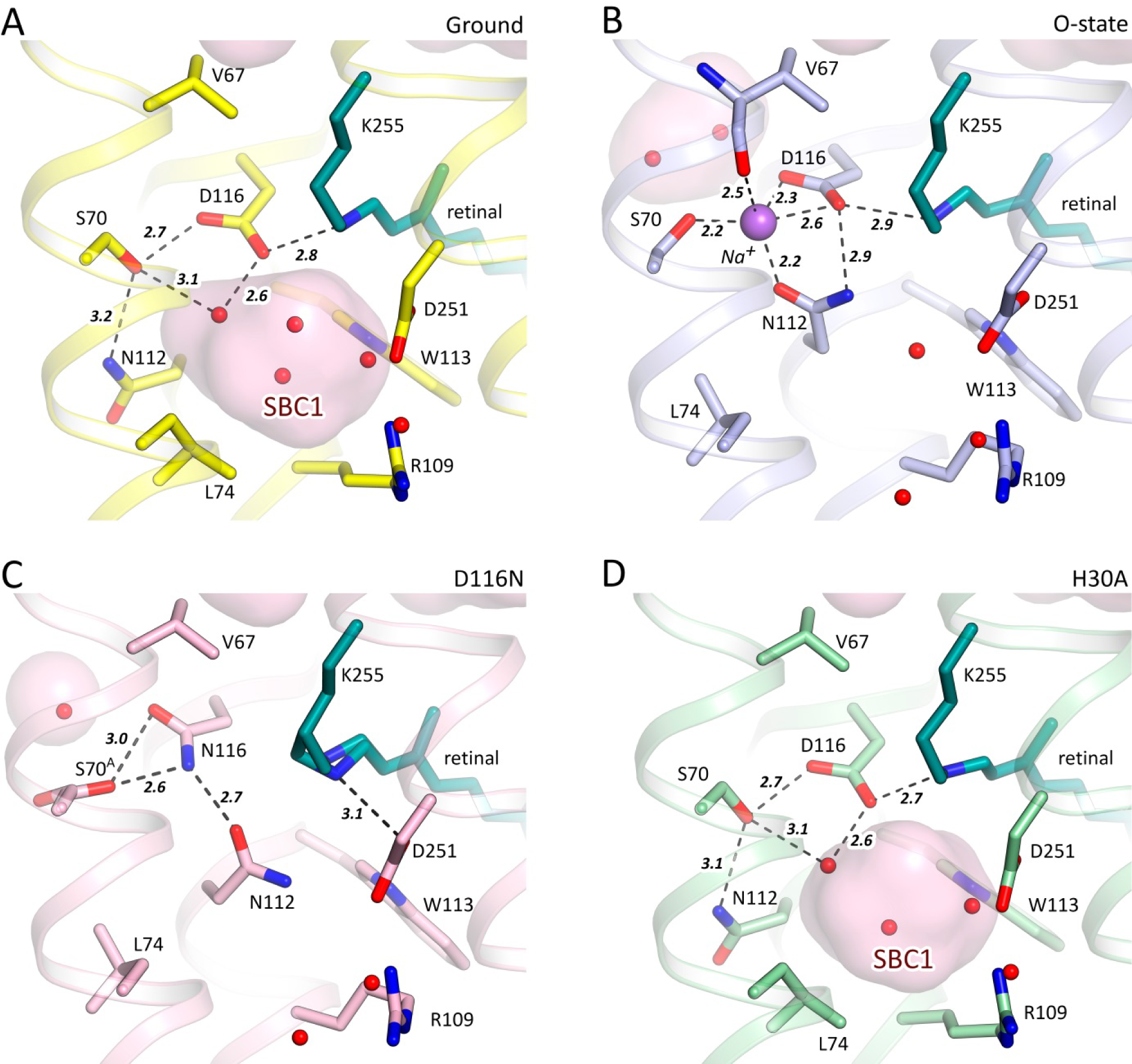
RSB region and Na^+^ binding site of KR2. **A.** The ground state of WT KR2 at pH 8.0 (PDB ID 6REW). **B.** The O-state of KR2 at pH 8.0 (present work). **C.** D116N mutant. **D.** H30A mutant. Cavities are calculated using HOLLOW^49^ shown in pink and marked with red labels. Retinal cofactor is colored teal. Water molecules are shown with red spheres. Sodium ion is shown with a purple sphere. Hydrogen bonds involving S70, N112, D116, D251 and RSB are shown with black dashed lines. The lengths of the shown hydrogen bonds are shown with bold italic numbers and are given in Å. Helix A and SBC2 are hidden for clarity.

**Supplementary Figure 14.**
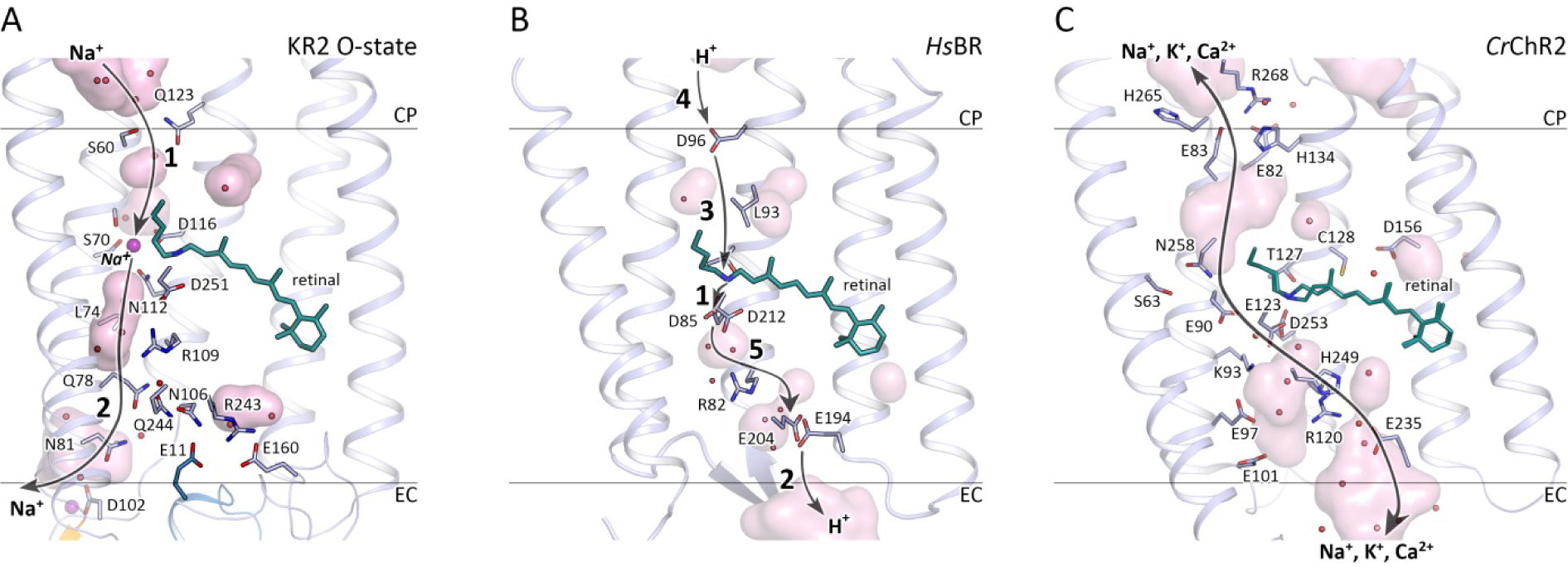
Comparison of ion pathways in different classes of microbial rhodopsins. **A.** The O-state of the KR2 (present work). **B.** The ground state of the *Hs*BR^57^ (PDB ID: 1C3W). **C.** Ground state of the *Cr*ChR2^27^ (PDB ID: 6EID). Cavities are calculated using HOLLOW^49^ and shown with pink surface. Gray lines indicate membrane hydrophobic/hydrophilic boundaries. Gray arrows indicate ion translocation pathways. In case of KR2 1 is for the Na+ uptake and 2 is for Na+ release. In case of *Hs*BR 1 is for proton translocation from the RSB to D85, 2 is for proton release from E194-E204 pair to extracellular space, 3 is for proton translocation from D96 to the RSB, 4 is for the D96 reprotonation from the cytoplasmic space and 5 is for the proton relocation from D85 to E194-E204 pair. In case of *Cr*ChR2 gray arrow indicate the ion pathway through the channel.

**Supplementary Figure 15.**
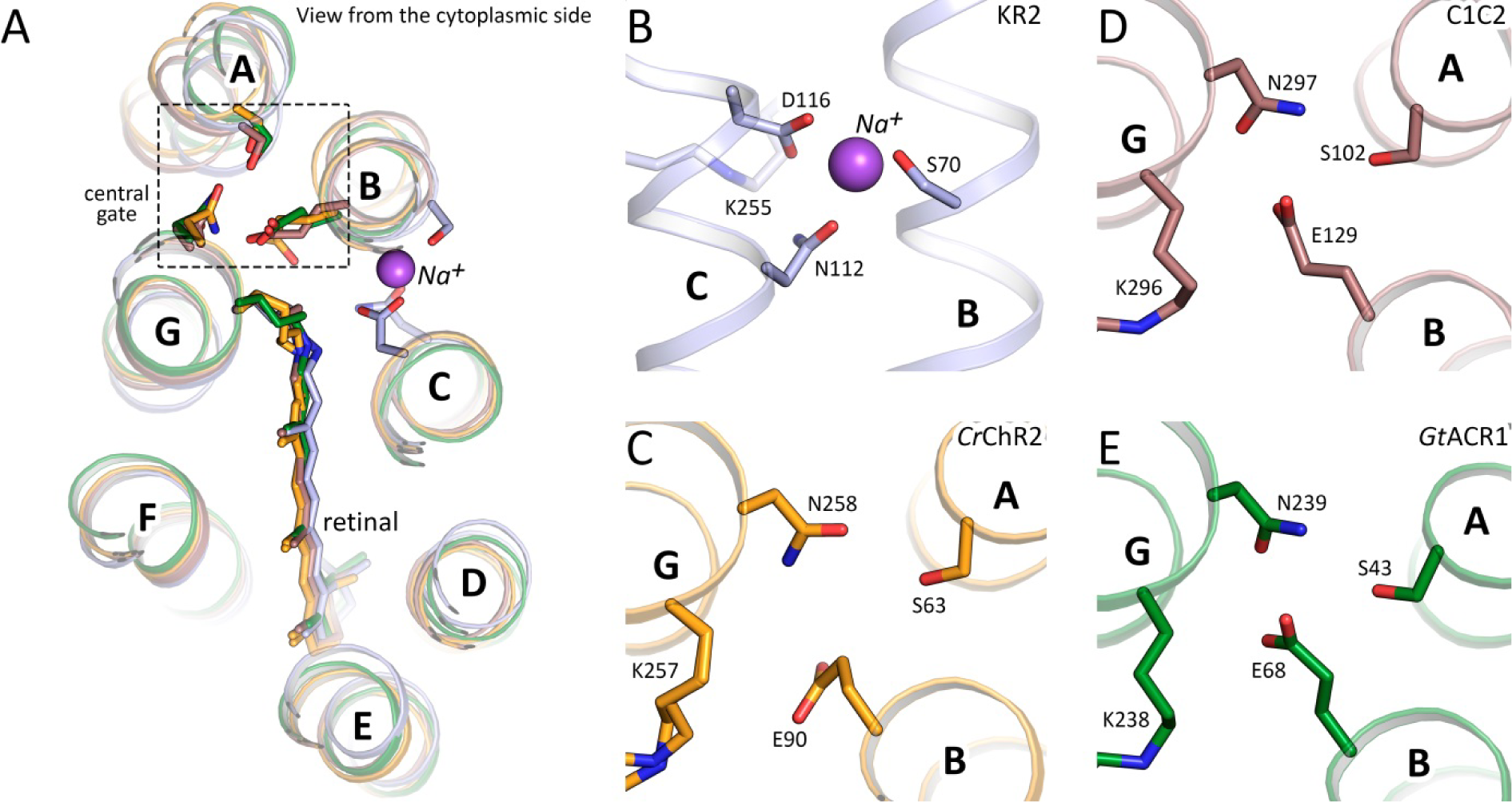
Na^+^ binding site of KR2 and central gates of channelrhodopsins. **A.** Structural alignment of KR2 (present work, blue), *Cr*ChR2 (PDB ID: 6EID, orange), C1C2 (PDB ID: 3UG9, brown) and *Gt*ACR1 (PDB ID: 6CSM, green). **B.** Transient sodium binding site in KR2. **C.** Central gate of *Cr*ChR2. **D.** Central gate of C1C2. **E.** Central gate of *Gt*ACR1.

**Supplementary Figure 16.**
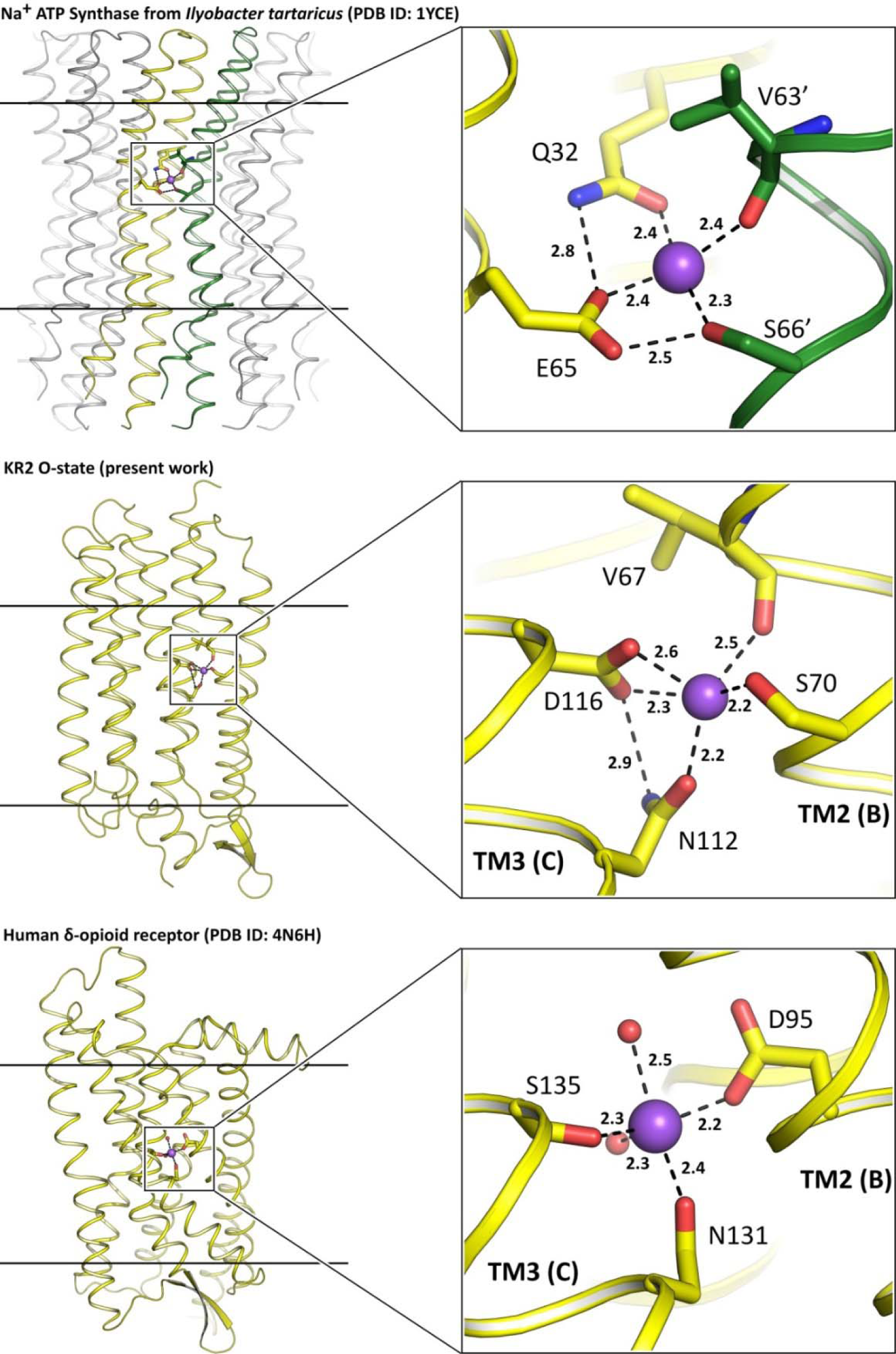
Na^+^ binding sites of different families of membrane proteins.

**Supplementary Figure 17.**
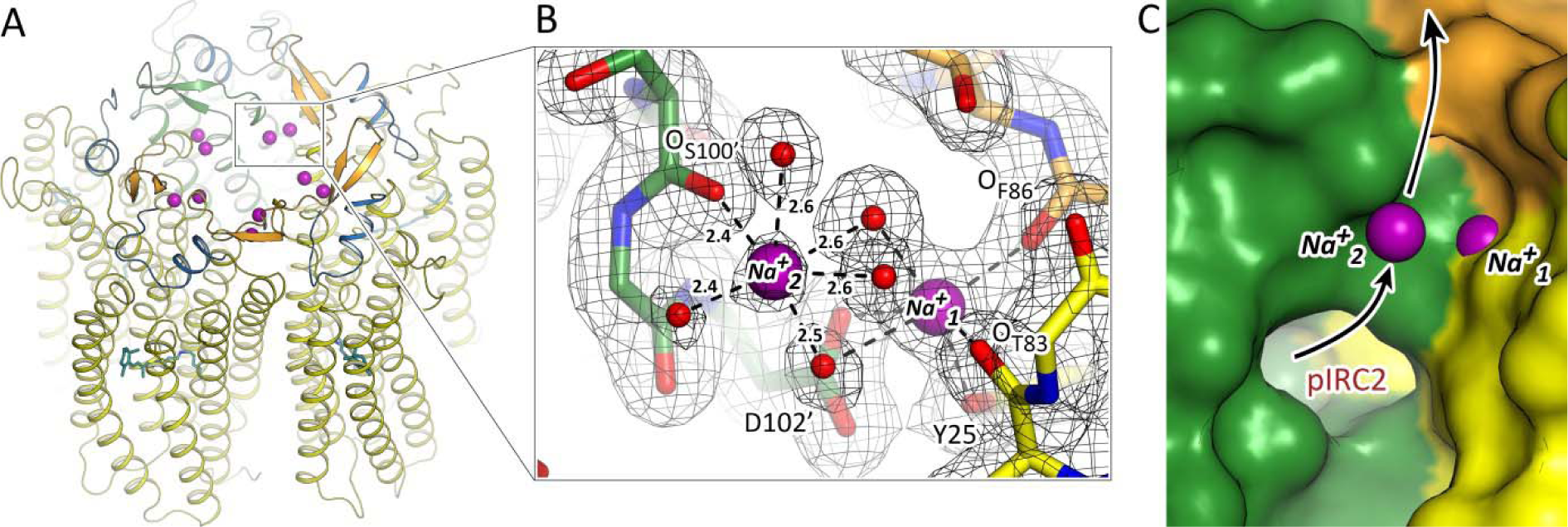
Additional Na^+^ identified at the KR2 interface in the ground state. **A.** Overall view of KR2 pentamer in the ground state from the extracellular side. Na^+^ bound at the protein surface are shown with purple spheres. **B.** Zoomed-in view of the Na^+^- binding sites. 2F_o_-F_c_ electron density maps around the Na^+^ and interacting residues and water molecules are shown with black mesh and are contoured at the level of 1.2 σ Distances between Na^+^ and nearby oxygens are shown with black dashed lines. Distances are in Å. C. Surface of KR2 pentamer near Na^+^. Putative ion-release cavity 2 (pIRC2) is labeled. Black arrows indicate putative relay pathway of the Na^+^ release.

**Supplementary Figure 18.**
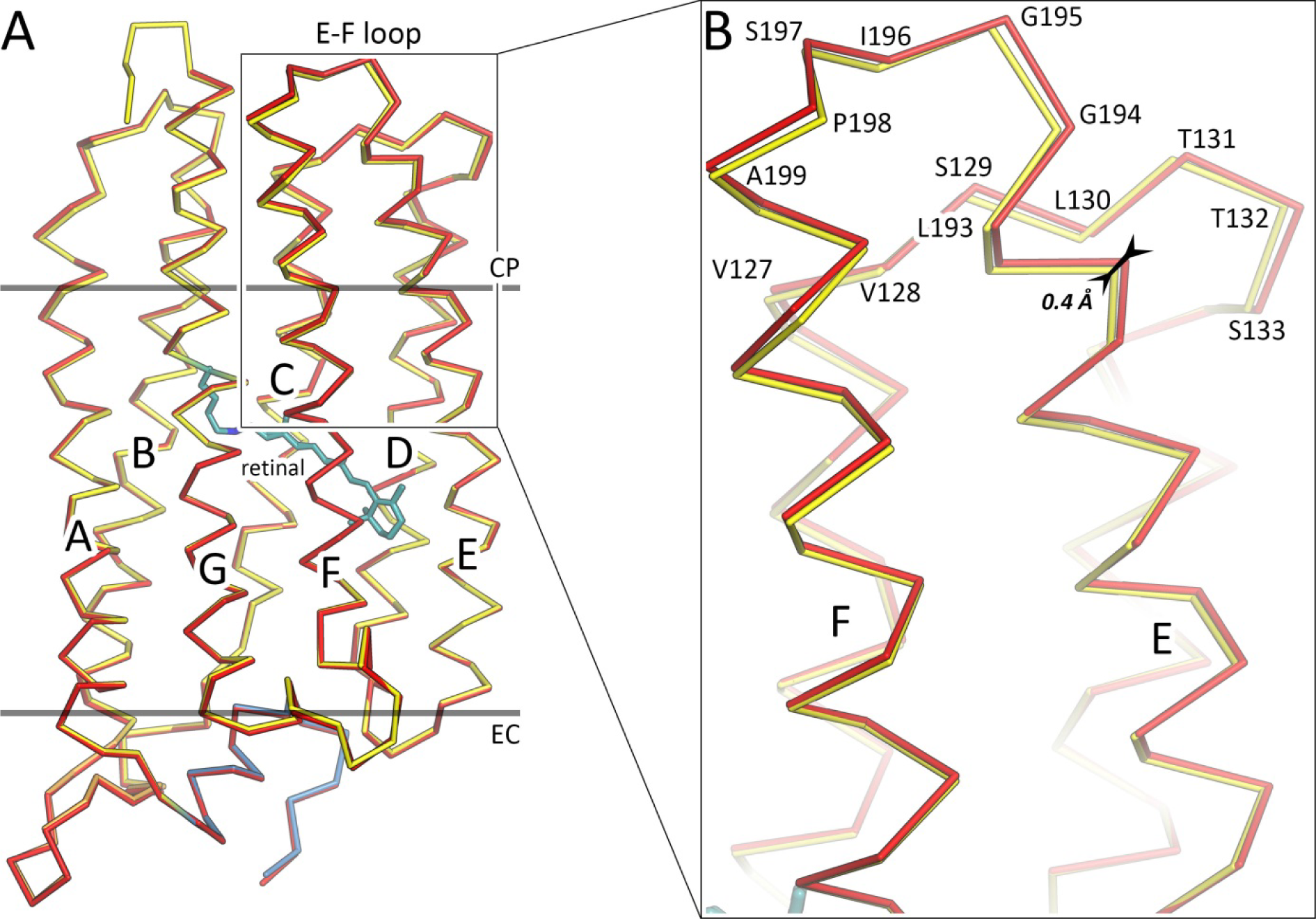
Structural alignment of KR2 protomers at 100K and 293K. **A.** Overall alignment. N-terminal α helix and N terminus are colored blue. BC loop, containing the β sheet, is colored orange. Membrane core boundaries are shown with black lines. KR2 ground state model at 100 and 293K are colored yellow and red, respectively. Retinal cofactor is colored teal. **B.** Enlarged view of the most notable rearrangements in protomer backbone. Helices are indicated with bold capital letters. Residues comprising the E-F and C-D loops are labeled. Shift of the helix E is shown with the black arrow and the distance is indicated.

**Supplementary Figure 19.**
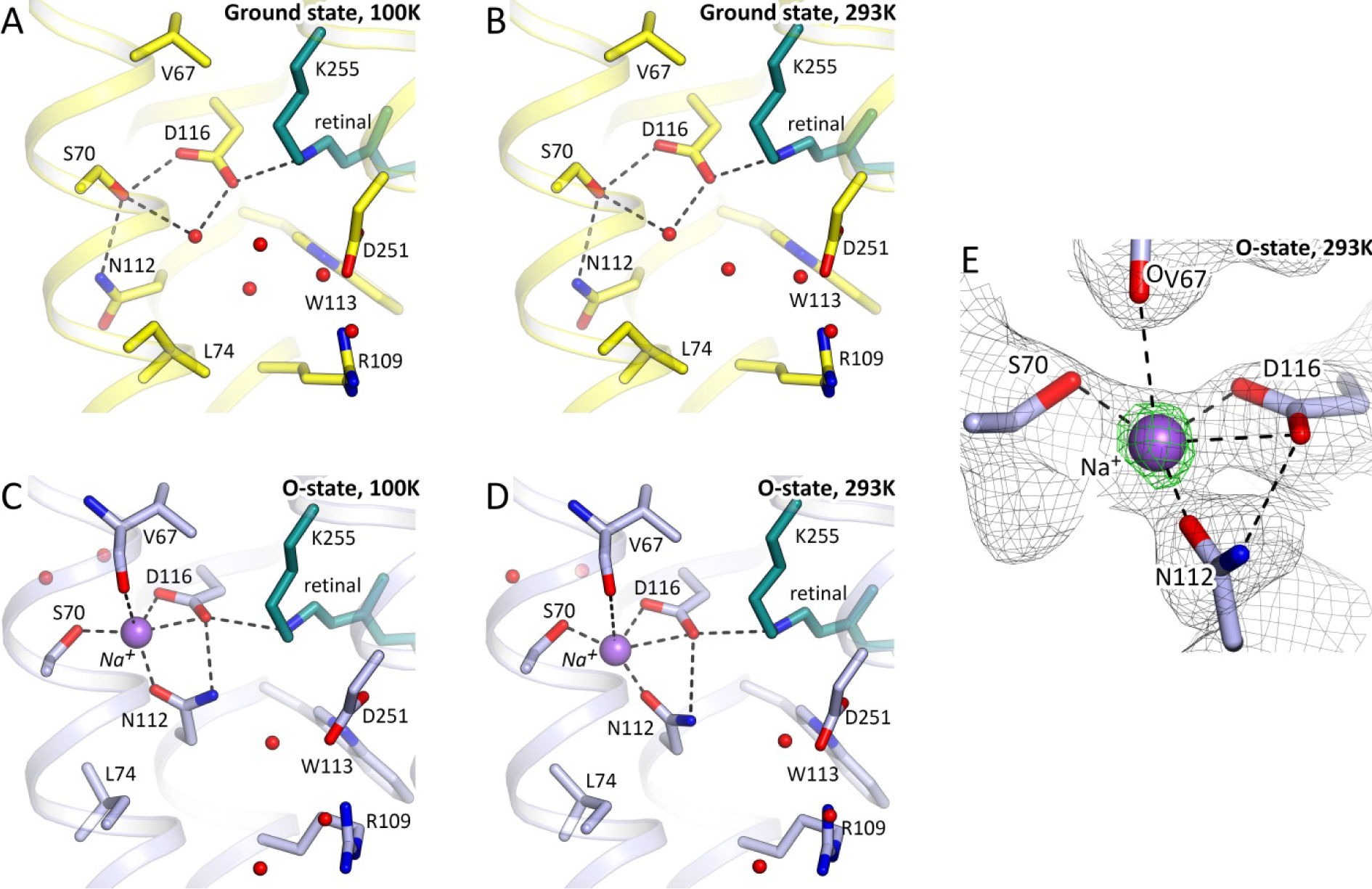
RSB region of the ground and the O-states of KR2 at 100K and 293K. **A.** The ground state of KR2 at 100K (PDB ID: 6REW). **B.** The ground state of KR2 at 293K (present work). **C.** The O-state of KR2 at 100K (present work). **D.** The O-state of KR2 at 293K (present work). Retinal cofactor is colored teal. Water molecules are shown with red spheres. Na^+^ is shown with a purple sphere. Hydrogen bonds are shown with black dashed lines. Helix A is hidden for clarity. **E.** Electron density maps (black, 2Fo-Fc at the level of 1.0σgreen, polder^20^ difference maps omitting Na^+^ contoured at the level of 4.0 the O-state at 293K.) of the Na^+^ binding site in the O-state at 293K.

**Supplementary Figure 20.**
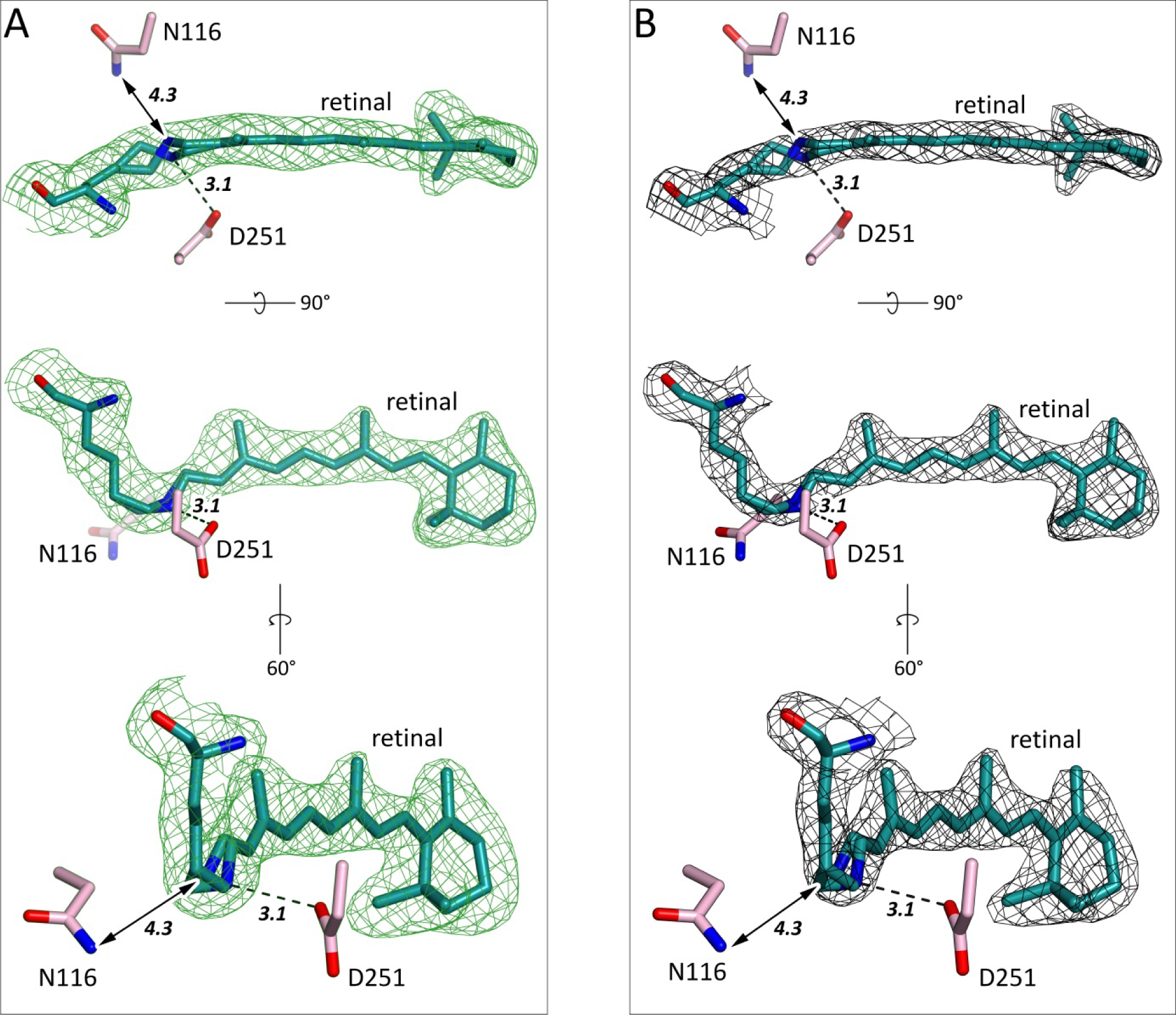
Examples of electron densities of the KR2-D116N. **A.** Polder^20^ maps for K255 and retinal cofactors of all five protomers of KR2-D116N structure. Maps are contoured at the level of 4.0 σ contoured at the level of 1.5 σ 2F_o_-F_c_ electron density maps around K255 and retinal cofactor Retinal and K255 are colored teal. Hydrogen bond between the.RSB and D251 is shown with black dashed lines. The distance between the RSB and nearest atom of N116 is shown with black arrowed line. The lengths of the hydrogen bond and the distance are shown with bold italic numbers and are in Å.

**Supplementary Figure 21.**
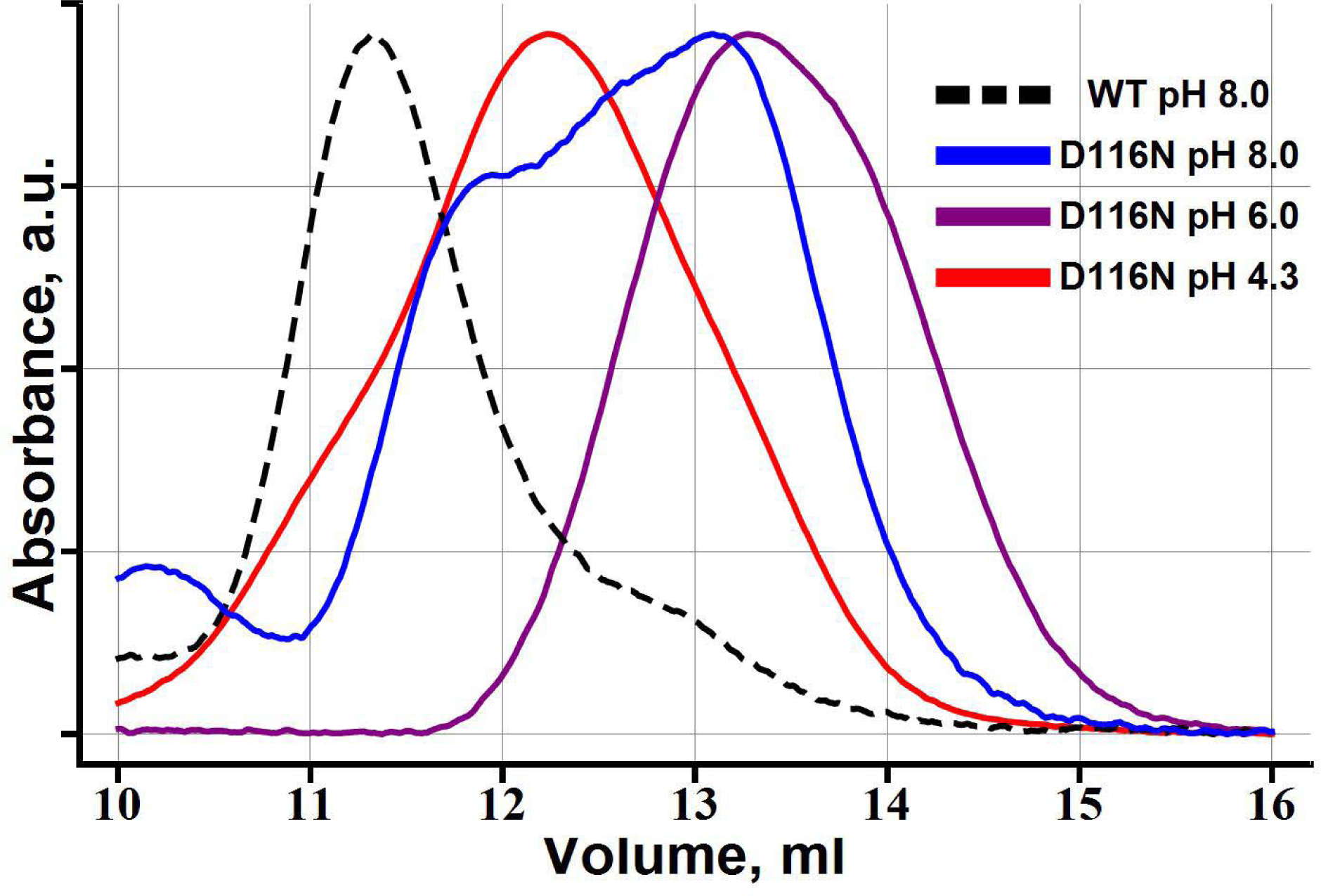
Size exclusion chromatography profiles of D116N mutant of KR2. Protein with initial concentration of 70 mg/ml was dissolved in buffer solution containing 200 mM NaCl with 0.1% DDM to final concentration of 1 mg/ml and dialyzed against 100x volume of the buffer of needed pH with no less than 5 times substitution of the outer buffer solution during at least 72 hours.

**Supplementary Figure 22.**
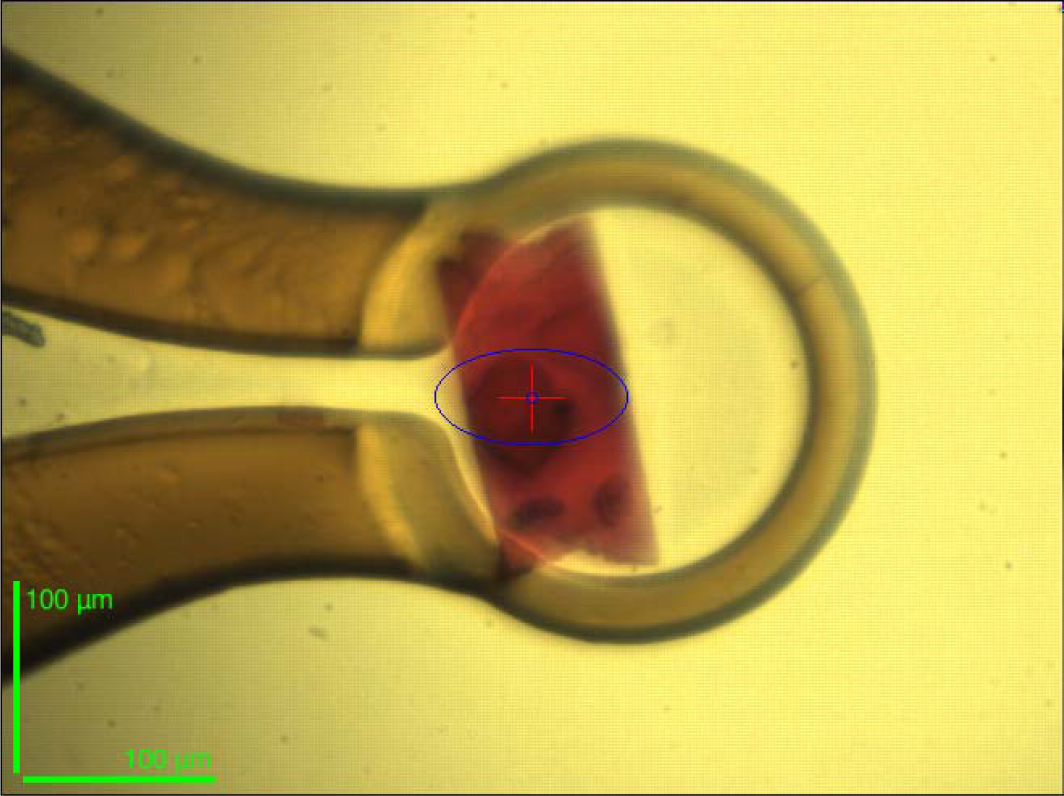
Example of KR2 crystals used in present work. The mean size of the crystal is 200x100x30 μm^3^

### Supplementary Videos

Supplementary Video 1. Exemplary trajectory of Na^+^ release obtained using molecular dynamics.

## References

1. Gushchin, I. & Gordeliy, V. Microbial Rhodopsins. in 19–56 (2018). doi:10.1007/978-981-10-7757-9_2

2. Inoue, K. et al. A light-driven sodium ion pump in marine bacteria. Nat. Commun. 4, (2013).

3. Gushchin, I. et al. Crystal structure of a light-driven sodium pump. Nat. Struct. Mol. Biol. 22, 390–396 (2015).

4. Kovalev, K. et al. Structure and mechanisms of sodium-pumping KR2 rhodopsin. Sci. Adv. 5, eaav2671 (2019).

5. Kato, H. E. et al. Structural basis for Na + transport mechanism by a light-driven Na + pump. Nature 521, 48–53 (2015).

6. Kato, Y., Inoue, K. & Kandori, H. Kinetic Analysis of H+-Na+ Selectivity in a Light-Driven Na+-Pumping Rhodopsin. J. Phys. Chem. Lett. (2015). doi:10.1021/acs.jpclett.5b02371

7. Abe-Yoshizumi, R., Inoue, K., Kato, H. E., Nureki, O. & Kandori, H. Role of Asn112 in a Light-Driven Sodium Ion-Pumping Rhodopsin. Biochemistry 55, 5790–5797 (2016).

8. Shevchenko, V. et al. Sodium and engineered potassium light-driven pumps. in Optogenetics: From Neuronal Function to Mapping and Disease Biology (2017). doi:10.1017/9781107281875.008

9. Vogt, A. et al. Engineered Passive Potassium Conductance in the KR2 Sodium Pump. Biophys. J. (2019). doi:10.1016/j.bpj.2019.04.001

10. Shibata, M. et al. Oligomeric states of microbial rhodopsins determined by high-speed atomic force microscopy and circular dichroic spectroscopy. Sci. Rep. 8, (2018).

11. Gushchin, I. et al. Structure of the light-driven sodium pump KR2 and its implications for optogenetics. FEBS J. 283, 1232–1238 (2016).

12. Kaur, J. et al. Solid-state NMR analysis of the sodium pump Krokinobacter rhodopsin 2 and its H30A mutant. J. Struct. Biol. (2018). doi:10.1016/j.jsb.2018.06.001

13. Kandori, H., Inoue, K. & Tsunoda, S. P. Light-Driven Sodium-Pumping Rhodopsin: A New Concept of Active Transport. Chemical Reviews (2018). doi:10.1021/acs.chemrev.7b00548

14. Nishimura, N., Mizuno, M., Kandori, H. & Mizutani, Y. Distortion and a Strong Hydrogen Bond in the Retinal Chromophore Enable Sodium-Ion Transport by the Sodium-Ion Pump KR2. J. Phys. Chem. B (2019). doi:10.1021/acs.jpcb.9b00928

15. Gerwert, K., Freier, E. & Wolf, S. The role of protein-bound water molecules in microbial rhodopsins. Biochimica et Biophysica Acta - Bioenergetics (2014). doi:10.1016/j.bbabio.2013.09.006

16. Oesterhelt, D. & Stoeckenius, W. Rhodopsin-like protein from the purple membrane of Halobacterium halobium. Nat. New Biol. 233, 149–152 (1971).

17. Gordeliy, V. I. et al. Molecular basis of transmembrane signalling by sensory rhodopsin II–transducer complex. Nature 419, 484–487 (2002).

18. Moukhametzianov, R. et al. Development of the signal in sensory rhodopsin and its transfer to the cognate transducer. Nature 440, 115–119 (2006).

19. Weinert, T. et al. Proton uptake mechanism in bacteriorhodopsin captured by serial synchrotron crystallography. Science (80-.). (2019). doi:10.1126/science.aaw8634

20. Liebschner, D. et al. Polder maps: Improving OMIT maps by excluding bulk solvent. Acta Crystallogr. Sect. D Struct. Biol. (2017). doi:10.1107/S2059798316018210

21. Chen, H. F. et al. Time-resolved FTIR study of light-driven sodium pump rhodopsins. Phys. Chem. Chem. Phys. (2018). doi:10.1039/c8cp02599a

22. Balashov, S. P. et al. Light-driven Na+pump from Gillisia limnaea: A high-affinity Na+binding site is formed transiently in the photocycle. Biochemistry (2014). doi:10.1021/bi501064n

23. Yoshizawa, S. et al. Functional characterization of flavobacteria rhodopsins reveals a unique class of light-driven chloride pump in bacteria. Proc. Natl. Acad. Sci. U. S. A. (2014). doi:10.1073/pnas.1403051111

24. Yun, J. H. et al. Non-cryogenic structure of a chloride pump provides crucial clues to temperature-dependent channel transport efficiency. J. Biol. Chem. (2019). doi:10.1074/jbc.RA118.004038

25. Shigeta, A. et al. Long-distance perturbation on Schiff base–counterion interactions by His30 and the extracellular Na + -binding site in Krokinobacter rhodopsin 2. Phys. Chem. Chem. Phys. 20, 8450–8455 (2018).

26. Shevchenko, V. et al. Inward H^+^pump xenorhodopsin: Mechanism and alternative optogenetic approach. Sci. Adv. 3, (2017).

27. Volkov, O. et al. Structural insights into ion conduction by channelrhodopsin 2. Science (80-.). 358, (2017).

28. Studier, F. W. Protein production by auto-induction in high-density shaking cultures. Protein Expr. Purif. (2005). doi:10.1016/j.pep.2005.01.016

29. Landau, E. M. & Rosenbusch, J. P. Lipidic cubic phases: A novel concept for the crystallization of membrane proteins. Proc. Natl. Acad. Sci. 93, 14532–14535 (1996).

30. Caffrey, M. & Cherezov, V. Crystallizing membrane proteins using lipidic mesophases. Nat. Protoc. 4, 706–731 (2009).

31. Gushchin, I. et al. Structural insights into the proton pumping by unusual proteorhodopsin from nonmarine bacteria. Proc. Natl. Acad. Sci. 110, 12631–12636 (2013).

32. Gordeliy, V. I. et al. Molecular basis of transmembrane signalling by sensory rhodopsin II-transducer complex. Nature 419, 484–487 (2002).

33. Chizhov, I. & Engelhard, M. Temperature and halide dependence of the photocycle of halorhodopsin from Natronobacterium pharaonis. Biophys. J. 81, 1600–1612 (2001).

34. Chizhov, I. et al. The photophobic receptor from Natronobacterium pharaonis: Temperature and pH dependencies of the photocycle of sensory rhodopsin II. Biophys. J. 75, 999–1009 (1998).

35. Von Stetten, D. et al. In crystallo optical spectroscopy (icOS) as a complementary tool on the macromolecular crystallography beamlines of the ESRF. Acta Crystallogr. Sect. D Biol. Crystallogr. (2015). doi:10.1107/S139900471401517X

36. Kabsch, W. XDS. Acta Crystallogr D Biol Crystallogr (2010). doi:10.1107/S0907444909047337

37. Winn, M. D. et al. Overview of the *CCP* 4 suite and current developments. Acta Crystallogr. Sect. D Biol. Crystallogr. (2011). doi:10.1107/S0907444910045749

38. Tickle, I. J. et al. STARANISO. Cambridge, United Kingdom: Global Phasing Ltd. (2018).

39. Vagin, A. & Teplyakov, A. Molecular replacement with MOLREP. Acta Crystallogr. Sect. D Biol. Crystallogr. (2010). doi:10.1107/S0907444909042589

40. Murshudov, G. N. et al. REFMAC5 for the refinement of macromolecular crystal structures. Acta Crystallogr. D. Biol. Crystallogr. 67, 355–367 (2011).

41. Adams, P. D. et al. PHENIX: A comprehensive Python-based system for macromolecular structure solution. Acta Crystallogr. Sect. D Biol. Crystallogr. (2010). doi:10.1107/S0907444909052925

42. Emsley, P. & Cowtan, K. Coot: model-building tools for molecular graphics. Acta Crystallogr. Sect. D Biol. Crystallogr. 60, 2126–2132 (2004).

43. Jo, S., Kim, T., Iyer, V. G. & Im, W. CHARMM-GUI: A web-based graphical user interface for CHARMM. J. Comput. Chem. 29, 1859–1865 (2008).

44. Abraham, M. J. et al. GROMACS: High performance molecular simulations through multi-level parallelism from laptops to supercomputers. SoftwareX 1–2, 19–25 (2015).

45. Huang, J. et al. CHARMM36m: an improved force field for folded and intrinsically disordered proteins. Nat. Methods 14, 71–73 (2017).

46. Zhu, S., Brown, M. F. & Feller, S. E. Retinal Conformation Governs p K a of Protonated Schiff Base in Rhodopsin Activation. J. Am. Chem. Soc. 135, 9391–9398 (2013).

47. Laio, A. & Parrinello, M. Escaping free-energy minima. Proc. Natl. Acad. Sci. 99, 12562– 12566 (2002).

48. Tribello, G. A., Bonomi, M., Branduardi, D., Camilloni, C. & Bussi, G. PLUMED 2: New feathers for an old bird. Comput. Phys. Commun. 185, 604–613 (2014).

49. Ho, B. K. & Gruswitz, F. HOLLOW: Generating accurate representations of channel and interior surfaces in molecular structures. BMC Struct. Biol. 8, (2008).

50. Volkov, O. et al. Structural insights into ion conduction by channelrhodopsin 2. Science (80-.). 358, (2017).

51. Kato, H. E. et al. Crystal structure of the channelrhodopsin light-gated cation channel. Nature (2012). doi:10.1038/nature10870

52. Li, H. et al. Crystal structure of a natural light-gated anion channelrhodopsin. Elife (2019). doi:10.7554/eLife.41741

53. Kim, Y. S. et al. Crystal structure of the natural anion-conducting channelrhodopsin GtACR1. Nature (2018). doi:10.1038/s41586-018-0511-6

54. Wietek, J. et al. Conversion of channelrhodopsin into a light-gated chloride channel. Science (80-.). 344, 409–412 (2014).

55. Shalaeva, D. N., Galperin, M. Y. & Mulkidjanian, A. Y. Eukaryotic G protein-coupled receptors as descendants of prokaryotic sodium-translocating rhodopsins. Biol. Direct (2015). doi:10.1186/s13062-015-0091-4

56. Shigeta, A. et al. Solid-state nuclear magnetic resonance structural study of the retinal-binding pocket in sodium ion pump rhodopsin. Biochemistry (2017). doi:10.1021/acs.biochem.6b00999

57. Luecke, H., Schobert, B., Richter, H. T., Cartailler, J. P. & Lanyi, J. K. Structure of bacteriorhodopsin at 1.55 Å resolution. J. Mol. Biol. 291, 899–911 (1999).

