## Supplementary Table 1 for "Molecular mechanism of light-driven sodium pumping"

**Supplementary Table 1. Data collection and refinement statistics.**

| | *WT-100K Ground state* | *WT-100K O-state* | *WT-293K Dark state* | *WT-293K Illuminated state* | *D116N-100K monomeric* | *D116N-100K pentameric* | *H30A-100K pentameric* |
|---|---|---|---|---|---|---|---|
| pH | 8.0 | 8.0 | 8.0 | 8.0 | 4.6 | 8.0 | 8.0 |
| **Data collection** | | | | | | | |
| Space group | $C222_1$ | $C222_1$ | $C222_1$ | $C222_1$ | I222 | $C222_1$ | $C222_1$ |
| Number of crystals | 1 | 1 | 3 | 3 | 1 | 1 | 1 |
| *Cell dimensions* | | | | | | | |
| $a$, $b$, $c$ (Å) | 131.87, 240.32, 135.51 | 131.16, 240.63, 135.04 | 135.15, 239.89, 138.35 | 134.92, 239.78, 138.35 | 40.89, 83.60, 233.83 | 131.34, 240.48, 135.41 | 131.09, 239.73, 135.13 |
| $\alpha$, $\beta$, $\gamma$ (°) | 90, 90, 90 | 90, 90, 90 | 90, 90, 90 | 90, 90, 90 | 90, 90, 90 | 90, 90, 90 | 90, 90, 90 |
| Resolution (Å) | 48.16-2.00 (2.03-2.00) | 48.06-2.10 (2.14-2.10) | 48.84-2.50 (2.55-2.50) | 48.81-2.59 (2.65-2.59) | 40.81-1.80 (1.84-1.80) | 48.13-2.35 (2.39-2.35) | 48.01-2.20 (2.24-2.20) |
| $R_{merge}$ (%) | 7.6 (174.9) | 7.8 (276.0) | 21.9 (236.5) | 23.7 (213.1) | 4.6 (100.0) | 11.0 (192.5) | 12.2 (209.9) |
| $R_{pim}$ (%) | 3.0 (67.4) | 2.2 (75.7) | 5.4 (81.7) | 8.6 (77.1) | 2.0 (40.6) | 4.5 (80.8) | 4.9 (82.7) |
| $I/\sigma I$ | 13.2 (1.2) | 20.1 (1.1) | 10.1 (1.2) | 6.7 (1.1) | 16.2 (1.7) | 11.3 (1.0) | 11.5 (0.9) |
| *CC1/2 (%)* | 99.9 (74.3) | 99.8 (56.0) | 99.8 (81.7) | 99.4 (50.7) | 99.9 (90.5) | 99.9 (78.7) | 99.9 (48.9) |
| Completeness (%) | 99.9 (100.0) | 99.8 (99.9) | 100.0 (100.0) | 99.6 (94.8) | 99.8 (99.7) | 99.9 (99.9) | 98.9 (99.2) |
| Unique reflections | 144,508 (7135) | 123,865 (6133) | 78,134 (4418) | 69,594 (4259) | 37,771 (2191) | 89,161 (4530) | 106,450 (5261) |
| **Refinement** | | | | | | | |
| Resolution (Å) | 50-2.0 | 50-2.1 | 50-2.5 | 50-2.6 | 20-1.8 | 50-2.35 | 50-2.20 |
| No. reflections | 135,234 | 101,547 | 74,363 | 59,103 | 35,822 | 74,883 | 91,343 |
| $R_{work}/R_{free}$ (%) | 17.7/20.3 | 17.6/20.0 | 17.4/19.8 | 19.9/21.8 | 17.4/22.9 | 19.9/22.3 | 18.2/20.5 |
| *R.m.s deviations* | | | | | | | |
| Protein bond lengths (Å) | 0.0027 | 0.0027 | 0.0023 | 0.0012 | 0.0054 | 0.0096 | 0.0019 |
| Protein bond angles (°) | 1.0671 | 1.0479 | 1.1192 | 1.0765 | 0.7962 | 1.0876 | 1.0502 |
